# ClinCNV: multi-sample germline CNV detection in NGS data

**DOI:** 10.1101/2022.06.10.495642

**Authors:** German Demidov, Marc Sturm, Stephan Ossowski

## Abstract

Germline copy number variants (CNVs) are a common source of genomic variation involved in many genetic disorders, and their detection is crucial for clinical molecular diagnostics. Genomic microarrays, quantitative polymerase chain reaction (qPCR), and multiplex ligation-dependent probe amplification (MLPA) have been widely used for CNV detection in clinics for many years. Similarly, next-generation sequencing (NGS) applications such as whole-genome sequencing (WGS) and whole-exome sequencing (WES) are well-established, highly accurate techniques for the detection of single nucleotide variants (SNVs) and small insertions and deletions (indels). However, CNV detection using NGS remains challenging due to short read lengths, smaller than CNVs sizes. CNV detection using read coverage depths summarized in genomic regions is affected by various biases that arise during the library preparation and sequencing. We have developed a novel strategy for detecting CNVs, implemented in the tool ClinCNV (freely available on https://github.com/imgag/ClinCNV). ClinCNV does multi-sample normalization and CNV calling, using an original algorithm taking the best from the circular binary segmentation method and Hidden Markov model-based approaches. Here, we describe the methods and discuss the results obtained by applying ClinCNV to thousands of clinical WES, WGS, and shallow-WGS samples in various clinical and research settings.

## I. Introduction

Short read high throughput sequencing remains the predominantly applied tool for SNVs and indels calling in clinical diagnostics of genetic diseases. Despite the release of many NGS-based CNV callers, the detection of CNVs in clinical applications remains challenging. There are several reasons for the reluctance of clinicians to accept WES or even WGS-based CNV detection as a replacement for microarrays and MLPA or qPCR. Foremost, the low sensitivity and recall of previously published tools have been discussed in detail in a large number of benchmark papers on real-life (non-simulated) WES data (e.g. [Trost et al., 2018, Whitford et al., 2019, Yao et al., 2017, Roller et al., 2016, D’Aurizio et al., 2016]). Also, tools often lack crucial features, such as parent-child trio calling or means for adequate quality control and visualization required for efficient use of CNV callers in clinical practice.

In this study, we aimed to develop a tool for reliable detection of CNVs that fulfils clinical testing needs. It includes the possibility to work with different data types, such as WES, WGS and shallow-WGS, in different calling modes such as standard single sample, trio and multi-sample cohort analysis.

Using the developed tool called ClinCNV, we investigated the following questions.

First, we wanted to understand if the current gold standard tools for CNVs detection in clinics, e.g. SNP/CGH arrays and qPCR/MLPA, could reliably be replaced with WES, shallow WGS (2-7x coverage) or WGS (>30x coverage) short-read sequencing.

Second, we wanted to understand if the developed tool’s performance is comparable to or better than the existing solutions. We benchmarked our tool against various existing bioinformatics methods on different data types, focusing on the limitations of the developed and existing approaches.

Finally, we raised the question if the classification and labelling of calls as true and false positive CNVs are appropriate for clinical applications. Clinical applications might favour sensitivity over specificity to guarantee that causal variants are not overlooked. It is usually impossible to ensure a high CNV detection power while keeping the false discovery rate (FDR) reasonable due to the high noisiness of short-read NGS data, especially WES. Therefore, instead of simply filtering out variants that are likely to be false positives, we advocate for an approach of estimating the FDR for each CNV, using machine learning methods and various characteristics of variants as predictors. Subsequently, CNVs annotated with FDRs can be further evaluated, combining probabilities of variants to be real (technical quality) and clinical metadata, such as gene-phenotype matching, presence of compound heterozygous mutations, segregation in the family or population allele frequencies (e.g., [Engelhardt et al., 2017]). It allows interpreters to balance the available resources for variants evaluation and validation, unlike in traditional hard filtering of low-quality variants.

## II. Materials and methods

### i. Read-depth based identification of rare CNVs

Since the tool was supposed to work not only with whole-genome but also with targeted sequencing data (WES or gene panels), we concentrated on the read-depth signal. Worth to note that other signals (such as B-allele frequency) can easily be added to the calling algorithm but is more beneficial for somatic analysis ([Demidov et al., 2019]) and not for germline diagnostics. The main steps performed by ClinCNV are:

1. binning of the genome or exome (using windows of uniform length for WGS or coordinates of design enrichment regions for WES or overlapping windows of reasonable size, usually 120bp, if intra-target CNVs detection is desired);
2. reads counting for each bin;
3. within-sample normalization (GC-content, region-length based, variance stabilization using square root transformation);
4. finding the samples whose coverage profiles are the most similar to the sample of interest normalized coverage, in other words, clustering of samples with similar coverage profiles;
5. between-sample normalization (using medians of coverages across the clusters of similar samples);
6. estimation of parameters for statistical models describing different copy-number states (taking into account per-sample and per-region variability);
7. calculation of the matrix of likelihoods, where the number of rows is equal to the number of predefined copy-number states, and the number of columns is equal to the number of bins;
8. segmentation and calling;
9. annotation with various QC metrics and visualization.

Since the first (normalization) steps are analogous to many other read-depth callers published previously, they are described in detail in the Supplementary. The segmentation and calling procedure was previously described in bioRxiv preprint of the somatic part of ClinCNV [Demidov et al., 2019], nevertheless, we describe it in the text of this paper too. The common CNV calling procedure is described in [Fawcett et al., 2022]. Here we concentrate on rare (less than 2.5% of a cohort) germline CNVs calling only.

For rare germline CNVs, we decided to limit the predefined copy-number space from 0 to 8. Each allowed copy-number defines a separate statistical model. We assume that most of the samples are diploid in each particular region. We rely on the assumption of Poisson distributed read counts. It allows us to apply a square root transformation, which makes the transformed normalized read counts distribution more similar to Normal and stabilizes variances. We can note that since most samples are assumed to be diploid, there are usually not enough data points to estimate parameters for each statistical model other than diploid. However, the variance stabilization allows us to estimate the variance for the diploid model only, using robust estimators and treating non-diploid data points as outliers, and then assume that other copy-number models have the same variance. We use robust estimators of standard deviation and location (*Q*_*n*_ [Rousseeuw et al., 1993], and median). Thus, for each copy-numbers from 0 to 8 we know the expected location shift and dispersion, which allows us to calculate a Normal likelihood for each data point (normalized coverage of a particular region in a particular sample). For each sample, we create a matrix of log-likelihoods of each data point across the genome under different copy-number assumptions. The matrix dimensions are |*G*|×| *S*| where *S* is the set of defined states (0 to 8), and *G* is the set of windows in the genome where read depth counts were summarized. Then, we assume the baseline diploid state for rare CNVs detection in autosomes or females X chromosome or haploid state for sex chromosomes in males. We subtract the baseline state’s log-likelihoods from other alternative states’ log-likelihoods. For each region and each sample, we have a logarithm of likelihood ratios, which is positive when the normalized coverage supports the alternative state more than the baseline and negative in the opposite case. The goal now is to find a stretch of consecutive genomic windows where the evidence of alternative states is the biggest, i.e., the segment with the strongest evidence of a CNV, which means the biggest positive sum of log-likelihoods.

We find such segments for all the states and choose the state with the largest evidence of non-baseline copy-number, using maximum sub-array sum algorithm [Bentley et al., 1984]. Then we segment the genome into three parts: to the left and right of the segment and the segment itself. We repeat this procedure until the newly found segments show evidence of an alternative state less than the pre-defined threshold.

CNV detection in trios is performed similarly, but the set of states is considered as copynumbers from 0 to 8 for the parents’ alleles, all possible combinations for the child copynumber and copy-number from 0 to 8 for *de novo* CNVs in the child. The corresponding matrices of likelihoods (estimated from mothers, fathers, and probands samples) are summed up into one, representing the whole trio likelihood of different combinations of different CNVs in family members.

Closely located windows, even nonoverlapping, may show a significant correlation of coverages and thus inflate log-likelihood. We correct it, doing a re-calculation of the likelihoods of neighbouring windows, using their joint likelihood of normalized coverage depths.

### ii. Execution, post-processing and annotation of calls in routine diagnostics using MegSAP pipeline

Benchmarking of clinical samples was done using the ClinCNV 1.16, implemented as a part of a Medical Genetics Sequence Analysis Pipeline (megSAP, https://github.com/imgag/megSAP).

Since the number of control samples (i.e., sequenced with the same protocol) can be large in a clinic or other medical genetics facility, the default clustering procedure implemented in ClinCNV, becomes impractical. Thus, as the first step of CNV calling, we sub-select 200 samples with the most similar coverage profile to the sample we want to analyse. We precalculate the matrix of between-sample similarities (Pearson correlation between depth-of-coverage profiles, excluding common CNV regions and sex chromosomes).

We also added a specific mode for the clinical diagnostics of CNVs in our pipeline called “superRecall”. In a superRecall mode, all the variants, even with the minimum evidence (loglikelihood of 3) of CNV presence, are detected. By default, such low-confidence CNVs are hidden in clinical GUI. However, investigation of such low-confidence variants could be decisive for some diseases with specific phenotypes or for cases with clear pathogenic short variants in recessive genes.

Three default filters in MegSAP pipeline were used in this paper: “high-sensitivity” (all the results + superRecall variants), “default” (variants with log-likelihood bigger than 20, overlap with common CNVs regions smaller than 80%), “array-like” (CNV-size bigger than 35kbps and overlap with common CNVs regions smaller than 95%).

The annotation of the calls includes allele frequency of variants observed in our inhouse database of CNVs and intersections with several other databases. For annotations we use DECIPHER [Bragin et al., 2014] (expert-reviewed clinical synopses for syndromes associated with CNVs), ClinGen Dosage Sensitivity Map [Riggs et al., 2018] (catalogue of genes and regions which are dosage-sensitive and recommended for targeting in cytogenomic arrays), pathogenic CNVs from ClinVar [Landrum et al., 2018] and HGMD [Stenson et al., 2017]. *CnvGeneAnnotation* script adds a list of all affected genes (transcripts extended by 5000 bp in every direction) from our in-house database for each CNV. Additionally, it annotates the gnomAD [Karczewski et al., 2019] observed/expected (oe) score for each gene and which part of the gene is affected (complete gene, intronic/intergenic or exonic/splicing).

### iii. Comparison of ClinCNV and alternative methods in WGS research cohorts of PCAWG and 1000GP

#### iii.1 Analysis of high coverage WGS data from PCAWG study

We compared the performance of rare deletions site detection in ClinCNV and paired-end mapping (PEM) based tool DELLY (which also uses Read Depth as additional evidence) using the large cohort of 2833 WGS samples (39x coverage on average) from the PCAWG study [ICGC/TCGA PCAWG et al., 2020]. We compared deletion sites only since the paired-end mapping methods are usually limited to tandem duplications detection; thus, these methods are not comparable for the detection of duplications.

For the False Discovery Rate estimation, we used the Intensity Rank Sum annotation test (IRS, [Sudmant et al., 2015]), using the data from 787 samples from the cohort, analyzed with Affymetrix Genome-Wide Human SNP Array 6.0. Both callsets were adjusted and had FDR estimated as less than 5% using Random Forest train-test approach. For simplicity, we concentrated on autosomal variants only.

The samples were sequenced in different centres at different times, using different protocols. ClinCNV and DELLY utilized the same cohort of samples; however, due to different requirements for data quality DELLY dataset consisted of 2642 genomes, and the ClinCNV dataset con-sisted of 2471 samples. We ended up with a cohort of 2336 samples that passed QC filterings for both tools. DELLY’s callset was prepared according to the procedure described in the preprint [Waszak et al., 2019]. ClinCNV variants detected in different samples were also merged, using max(2KB, 90%) reciprocal overlap intersection criteria. All the variants with a length of at least 1kbps and the log-likelihood score of 15 or more were used for the merging. The resolution (window size) in PCAWG analysis was equal to 1kbps.

ClinCNV FDR-adjusted dataset of deletion sites for comparison with DELLY consisted of 1) 2003 common CNVs (>5% allele frequency) and longer than or equal to 3 Kbps, 2) 20.084 deletion sites detected with rare CNV detection algorithm. Since the primary goal of ClinCNV was the detection of clinically relevant CNVs, we decided to concentrate on rare deletions in this comparison. Thus, only CNVs with a frequency of less than 5% in the studied cohort were evaluated. Additional filtering on DELLY variants suitable for comparison was performed. Only deletions that affected at least 500bp (half) in one of the 1 kbp windows were allowed since such a window would look more like copy-number neutral otherwise. Such filtering ended up in 19.518 deletion sites detected by ClinCNV and 24.893 deletion sites detected by DELLY.

The criteria for comparison used was: if the overlap between a deletion site detected by both ClinCNV and DELLY was bigger than the maximum of 75% of the longest variant length or 500 base pairs, these variants were considered the same.

#### iii.2 Comparison with 1000GP callset

An original structual variant callset from 3rd phase 1000 genomes project was used for the comparison [Sudmant et al., 2015].

### iv. Comparison of array-based and NGS-based method ClinCNV in a clinical setting

One of the main questions we had to answer was whether WES/WGS analysis with ClinCNV could replace arrays for CNV detection in clinical practice. More specifically, we aimed to compare the performance of ClinCNV as an NGS-based method with a conventional array approach in detecting CNVs larger than 50Kbps. The threshold of 50Kbps was chosen according to “Einheitlicher Bewertungsmassstab”, 1. Quartal of 2019, Paragraph 11508 – i.e. the German national guidelines on what is covered by the statutory health insurance, where is stated that only CNV detection methods with a resolution of 50 kilobases or better are allowed for postnatal total genomic examination of constitutional imbalances and only under condition that karyotyping using optical microscopy methods did not provide sufficient results.

Here and below, we use an acronym for Agilent SureSelect Human All Exon panels as ssHAE. The number after the acronym denotes the version of the panel.

All the in-house samples for which array intensities and sequencing results were available were selected as test samples. NGS results were WGS (TrueSeq PCR-free) and WES (ssHAE V6 and V7). The same samples were analyzed with the CytoScan platform (HD, 750K and Optima). Overall numbers of samples analysed with each platform are provided in table 1 and table 2. Only high-quality samples were used, resulting in less than 300 CNVs detected with the array-based method. All the arrays were analyzed with the tool provided by the array’s manufacturer, according to the guidelines.

**Table 1:** Overview of NGS analysed samples available

| NGS platform | Num of samples |
| --- | --- |
| WGS TruSeq PCR-free | 9 |
| WES ssHAEv6 | 197 |
| WES ssHAEv7 | 79 |
| Overall: | 285 |

**Table 2:** Overview of array analysed samples available

| Microarray platform | Num of samples |
| --- | --- |
| CytoScan HD | 39 |
| CytoScan 750K | 217 |
| CytoScan Optima | 12 |
| Overall: | 268 |

### v. Sensitivity of ClinCNV for detection of CNVs longer than 15kbp in shallow WGS data

One of the questions we wanted to answer is to which extent CNV detection can be done using shallow whole-genome sequencing. A cohort of 65 samples with known clinically relevant CNVs detected by alternative methods such as SNP microarrays, MLPA or high-depth whole-exome sequencing was formed.

Initially, shallow sequencing was planned to be around 4x depth. Nonetheless, due to the significant non-uniformity of library preparations, samples were sequenced with depth from 0.3x to 9.0x. After estimating the potential level of signal-to-noise ratio, we decided to calculate the coverage depth of such samples for windows of 5 kbps and aim to detect CNVs longer than three consecutive windows.

### vi. Detection of CNVs in WES data

#### vi.1 Comparison between ClinCNV and ExomeDepth using research WES samples

ExomeDepth [Plagnol et al., 2012] is a versatile tool that was used multiple times in different studies for the germline calling of CNVs in whole-exome data. The study itself was published in 2012, but the tool was updated multiple times and, according to the manual, the last update was done in August 2019. ExomeDepth v. 1.1.12 was used for comparison.

We had 40 samples from the CLL study [Puente et al., 2015] sequenced with ssHAE v.5 kit with BAM files available, and we decided to test ClinCNV and ExomeDepth using these 40 samples. The sequencing depth was relatively low, around 40x (Suppl. Figure 9). For the comparison, we used Intensity Rank Sum test, as described before. FDR was estimated as two multiplied by the number of p-values bigger than 0.5, divided by the overall number of CNVs.

#### vi.2 Comparison between ClinCNV and ExomeDepth using well covered clinical WES samples

Since the samples we used for comparison were covered with fewer reads than is expected by modern standards, we decided to perform an additional check using 70 in-house clinical exomes (coverage around 130x), sequenced with ssHAE v7 panel. These samples were also analysed with CytoScan 750K arrays. We used ClinCNV results obtained from the clinical routine. However, we tested ExomeDepth only within these 70 samples since our dataset consisted of more than 4.500 exomes, and it was not possible to analyse them all with ExomeDepth in a feasible time. CNVs were post-processed similarly - CNVs with less than 10% of the length or one kbps distance between breakpoints were merged. Then we post-filtered results of ClinCNV to have approximately the same FDRs as ExomeDepth, using the simple strategy (hard thresholds for overall log-likelihood and likelihood per target region). It was not possible to achieve exactly the same numbers due to rounding up the quality values of variants.

#### vi.3 Detection of CNVs in clinical WES data in trios

The algorithm for detection of variants in trios was applied to two available cohorts of WES sequenced samples – 634 samples were sequenced with ssHAEv6 exome enrichment, and 317 were sequenced with ssHAEv7, with a total of 951 samples. Interestingly we did not have only 317 trios (951/3), but 332. The reason is that in some families, several siblings were sequenced, and their parents’ DNA samples were sequenced only once (i.e. these are not trios, but quadruples or larger families, which we split into trios for this benchmark).

We concentrated on analysing autosomal CNVs since the interpretation of CNVs in sex chromosomes is complex. To assess the quality of a call, we have chosen the following rule. If a region in the trio was detected as copy number 2 in both parents and copy-number different than 2 in the child, we would consider this as a *de novo* event. We have tried different quality thresholds for called CNVs (20, 40, 60 and 80), taking into account that 20 is our default threshold for calling in WES samples.

In the beginning, we filtered out all the events that had more than 50% overlap with polymorphic regions detected in the PCAWG cohort. Since the density of enrichment regions of WES is uneven, we used not the reciprocal overlap but the overlap of 50% of the length of a detected variant as a measure of overlap. We used the same set of parameters we use for routine calling: the minimum length of a detected variant was set to 1 region, and clustering was performed, requiring at least 50 samples in each cluster. We filtered out samples that had more than 100 *de novo* CNVs or more than 1.500 CNVs detected in the whole trio of samples, ending up with a cohort of 235 families.

### vii. Estimation of quality of WES variants using array data and Quantile Random Forest Regression

Even though several thousand samples were sequenced with different exome enrichment kits in our clinics, we had no orthogonal microarray data for most of them. Therefore, we used only 255 samples that were analysed with microarrays and WES. (ssHAEv6, ssHAEv7 enrichment kits) Nevertheless, the whole cohort of samples was used for the normalisation (each sample was analysed with 200 most similar ones, as described above in MegSAP pipeline paragraph). The average depth of diagnostic exome samples was around 130x.

We have tried two modes of CNV detection, merging and validation: using all the data and using QC-filtered data and concluded that adding samples with many CNVs improves neither the predictive model nor the final quality of predictions. Thus, we excluded samples with more than 1750 CNVs called with a very high sensitivity: log-likelihood score of at least five and the intersection with common CNVs regions smaller than 80%. In the table 3 we describe our dataset. Overall, 235 out of 255 samples passed the QC control on the number of CNVs.

**Table 3:**
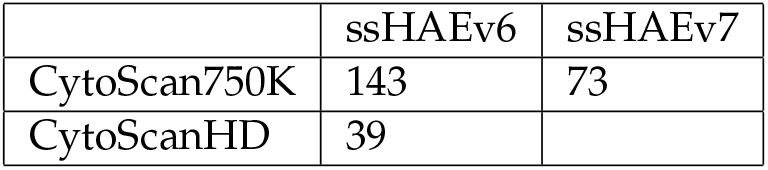
Number of samples available for testing.

|  | ssHAEv6 | ssHAEv7 |
| --- | --- | --- |
| CytoScan750K | 143 | 73 |
| CytoScanHD | 39 |  |

We use the same Intensity Rank Sum test as before to annotate our CNVs with p-values of the Wilxocon test (if at least one array probe is located within this CNV). Our goal for all CNVs was to estimate the probability of being a False Positive (FP) discovery (false-discovery rate, FDR). We tested our prediction procedure in 2 ways: within one kit and between kits. For ssHAEv6 kit we used 40% to 60% test-train set split. We also predicted FDR for CNVs in samples enriched with ssHAEv7 using 100% of variants detected in samples sequenced with ssHAEv6 kit as a train set.

We merged CNVs from different samples using 1kb difference in borders as the merging criterion. We kept our approach similar to the one used in IRS test. Since we intentionally included many FP CNVs, we could not separate True Positives (TP) and FPs simply by relying on the fact that p-values of FP variants are distributed uniformly, and p-value corresponding to TP variants are small. Instead of classifying variants as TP or FP, we decided to predict the expected p-value (which is closely related to FDR) from the Wilcoxon test, with one group containing array intensities in samples with a CNV call and another with array intensities in samples without this call. We used Quantile Regression Random Forest with two thousand trees to predict p-values (predictors described in Supplementary). For each variant, we had a distribution of predicted p-values instead of one value, using a discrete set of quantiles (from 0.5 to 0.999 with the step of 0.001). We decided to assign an FDR of *α* to a variant if *α* is the smallest quantile where the predicted p-value is still smaller than 0.5 and the next quantile is already bigger.

Thus, all the CNVs had a value assigned to them, from 0.002 to 1, which is double the FDR estimated at the previous step. We denote this value for the CNV *c* with *i*-th index *c*_*i*_ as *f dr*_*i*_. The real p-value obtained from the IRS test for the CNV will be denoted as *p*_*i*_. All the CNVs discovered will form a set denoted as *C*.

Thus, we may perform the following procedure for the test cohort. At first, we estimate the number of False Positive results as *FP* = 2 ·|*p* > 0.5| - 2 times the number of p-values greater than 0.5. Then, for FDR *α* we can find the following values:

- number of CNVs that have FDR less than *α*: |*C*_*α*_|, ∀*c*_*i*_∈ *C* : *f dr*_*i*_ < *α*
- estimated number of False Positive CNVs for all variants with estimated 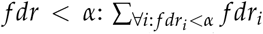:
- real number of False Positive CNVs for all variants with estimated *f dr* smaller then *α*: *FP*_*α*_ = 2×|*p*_*i*_ > 0.5 |,∀*i* : *f dr*_*i*_ < *α*;
- amount of True Positive events discovered: 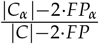 where for all CNVs *c*_*i*_ from *C*_*α*_ *f dr*_*i*_ < *α* (note: this value may be bigger than 1).

## III. Results

### i. Comparison of ClinCNV and alternative methods in WGS research cohorts of PCAWG and 1000GP

#### i.1 Comparison of ClinCNV and DELLY using PCAWG cohort

12.250 sites from DELLY generated callset were mapped to 12.189 sites from ClinCNV. Since the numbers are comparable, it shows that ClinCNV variants were not over-segmented. These numbers did not change a lot when more relaxed criteria for mapping were used (50% reciprocal overlap). ClinCNV detected 7.329 sites, missing in DELLY results and DELLY detected 12.643 CNVs not detected with ClinCNV.

Analyzing the histograms of variants obtained from different methods (fig. 1b), first of which (DELLY) uses paired-end distance and orientation information for CNVs detection and read-depth signature for filtering and genotyping. The second (ClinCNV) uses readdepth signature only. We conclude that:

1. ClinCNV and Delly detected many CNVs unique for a particular caller. Thus, both strategies should be applied for the detection of CNVs in WGS samples;
2. ClinCNV’s sensitivity breaks down for deletions that span less than three kbps, but it finds more unique events longer than ten kbps than DELLY;
3. Read-depth methods can detect different genomic duplications while paired-end methods are mainly limited to tandem duplications (comparison is not shown due to the absence of duplications in the DELLY generated dataset).

7.553 duplications and 20.084 deletions overall were detected in 2.471 WGS sequenced samples by ClinCNV (lengths are shown at fig. 2). Since the common CNVs were detected as a separate step, we checked if all the common CNVs were detected. The number of CNVs more frequent than 2.5% and detected via rare CNV detection algorithm was equal to 59 for deletions and 12 duplications. These reasonably small numbers show the efficiency of the common CNVs detection algorithm.

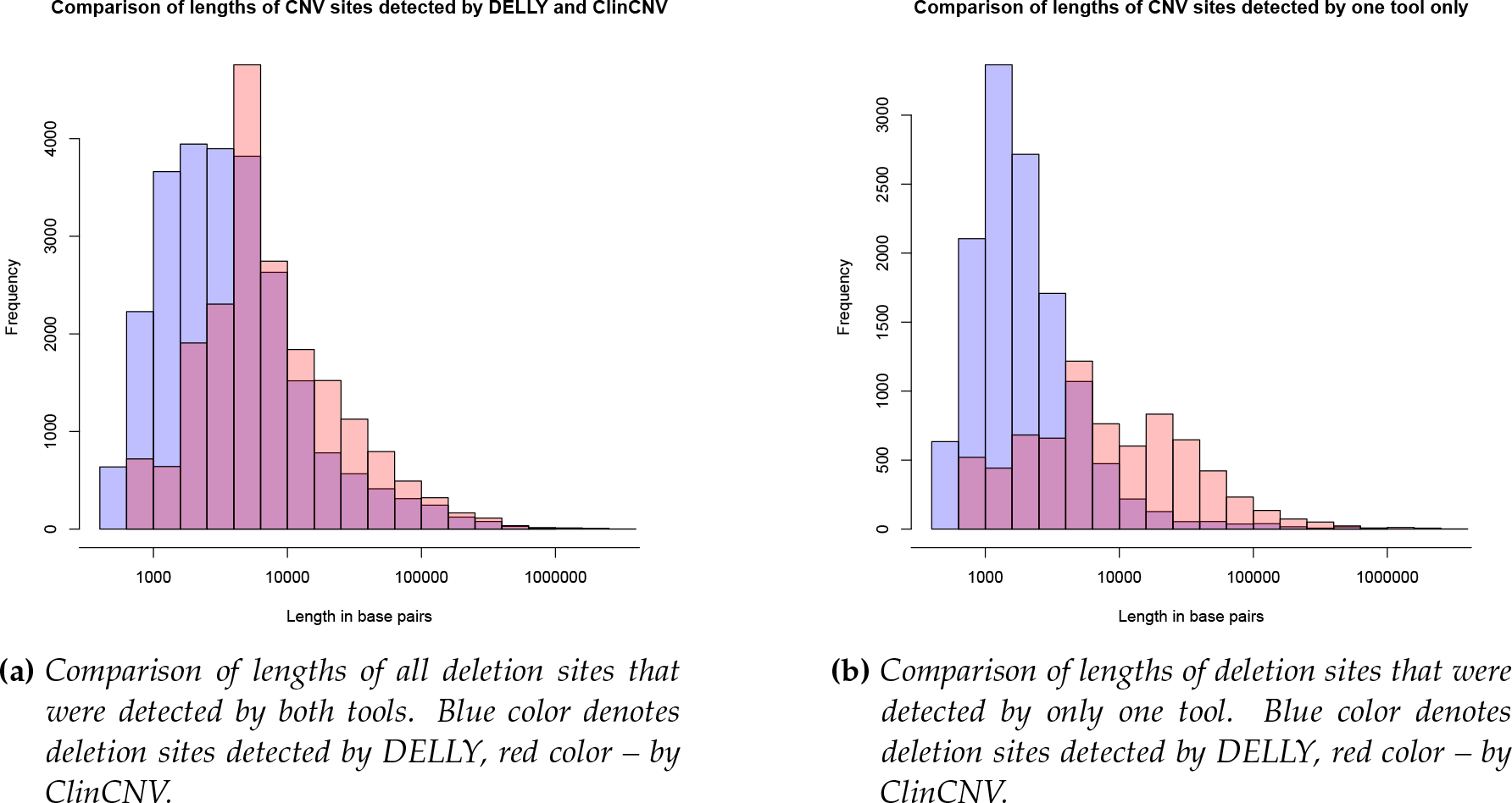

**Figure 2:**
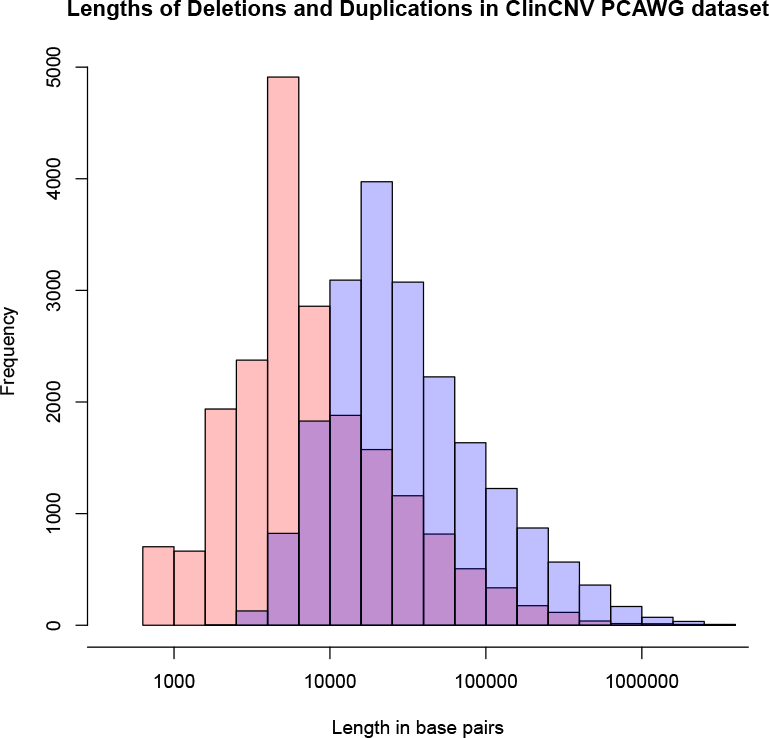
Lengths of detected sites. Red indicates deletions, blue duplications.

#### i.2 Comparison with 1000GP callset

Then we compared our callset with the callset of structural variants from the 1000 Genomes Project [Sudmant et al., 2015]. As before, we used ClinCNV’s callset of 7,553 rare duplication sites and 20,084 rare deletion sites for comparison with 35,868 deletion sites (and CNVs) and 8,954 duplication sites (and CNVs) longer than 500 base pairs from 1000GP callset. Only 5,682 of the ClinCNV deletions were presented in the 3rd phase 1000GP structural variant callset. For duplications, we used 50% overlap instead of 75% as a mapping criterion since the non-tandem duplications were detected using five kbps overlapping windows in 1000GP; thus, the resolution of our methods was different and less strict thresholds had to be applied. 1,392 rare duplications detected by ClinCNV mapped to 1,394 duplications (and CNVs) detected in the 3rd phase of 1000GP.

As a concluding remark for the analysis of PCAWG and 1000GP data, we can say that:

1. Read-depth method (ClinCNV) detects many deletion sites not presented in the callset of the paired-end method (DELLY) and vice versa, which means that neither method outperforms the other, and they have to be used together for the CNV calling;
2. 14,402 deletion sites and 6,161 duplication sites from our FDR-adjusted callset were not presented in the 1000GP 3rd phase SV callset, and thus, our callset can be considered a valuable source of information about structural variants in the human population. It is worth noting that the 1000GP analysis was done in a larger number of comparatively low covered whole genome samples, but many different methods were used for CNV calling. Thus, the power of detection was different.

### ii. Comparison between array-based and NGS-based method ClinCNV in a clinical setting

We concentrate on sensitivity since it is prioritized in clinical genetic diagnostics. Instead of the estimated specificity for clinical applications we provide an expected number of CNVs per sample (table 9) per different NGS method. It roughly (before disease-related gene filering) shows the actual burden of work a clinician can face. FDR of CNVs is described in details below (table 12).

#### ii.1 Sensitivity of ClinCNV in WGS data in detection of CNVs longer than 50 kilobases

We compared the callsets obtained using the default settings of ClinCNV (column name “Default”). A comparison using the High Sensitivity mode is provided under column name “All”.

Comparison of the ClinCNV callset obtained from high-coverage (> 30x) WGS samples was performed in a 9 sample cohort where only 22 suitable CNVs (longer than 50Kbps, not intersecting with polymorphic regions) were detected. Only one CNV was not found (table 4). It was covered by six array markers and was 57Kbps long. We found no visual evidence of such CNV in NGS coverage data, so we concluded that this CNV was a False Positive discovery of the array technology. Based on the high concordance between WGS-based and array-based results, we concluded that ClinCNV using WGS data could successfully replace array-based analysis for CNV detection in clinical diagnostics.

**Table 4:**
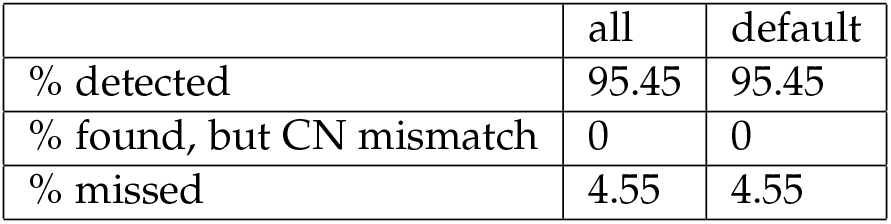
Results of comparison, ClinCNV in WGS and microarray detected CNVs, 22 CNVs discovered with microarray technology were used

#### ii.2 Sensitivity of ClinCNV for detection of CNVs longer than 50kbp in WES data

To compare the performance of CNV calling on whole-exome sequenced samples, we used ClinCNV in several different modes. First, we assessed the results with and without using off-target reads. Additionally, CNV calling on variants containing or not containing enrichment probes (e.g., purely intronic/intergenic and off-target) was assessed. Similarly to the previous analysis, we focused on CNVs that were longer than 50 kbps and had less than 20% intersection with CNPs. 406 CNVs called using array technology were used for validation.

ClinCNV missed 222 of 406 CNVs using default parameters (211 under high sensitivity mode). Missed CNVs were mainly located in intronic or intergenic regions that do not overlap with WES exons. For WES data without off-target reads we concluded that arrays could not be replaced with WES + ClinCNV analysis due to its inability to reliably detect CNVs in gene deserts, intergenic regions or long introns.

When ClinCNV calling quality was assessed on the CNVs overlapping with at least one enrichment probe, its performance improved.

Upon enforcing the intersection between CNVs and on-target probes, the amount of False Negative results dropped by more than 30%, as can be seen from table 6. Given the high number of False Negatives, we performed a manual check of the results.

**Table 5:** Comparison of array-based CNV calls with ClinCNV WES results (without off-target reads), 406 CNVs discovered with microarray technology were used

|  | all | default |
| --- | --- | --- |
| % detected | 45.07 | 43.35 |
| % found, but CN mismatch | 2.96 | 1.97 |
| % missed | 51.97 | 54.68 |

**Table 6:**
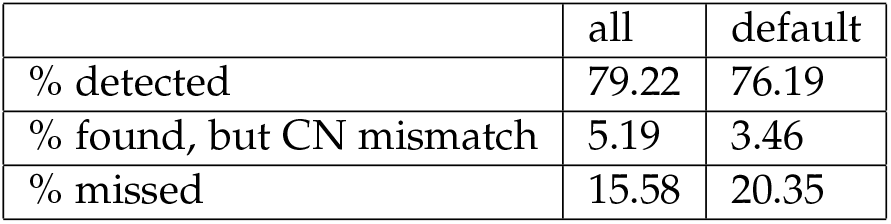
Comparison of array-based CNV calls with ClinCNV WES results (only CNVs containing at least one on-target probe), 192 CNVs detected in arrays were used.

|  | all | default |
| --- | --- | --- |
| % detected | 79.22 | 76.19 |
| % found, but CN mismatch | 5.19 | 3.46 |
| % missed | 15.58 | 20.35 |

As can be seen in table 7, we have found that slightly less than half of the False Negative variants in NGS are likely artefacts.

**Table 7:** Results of manual check of array-based CNVs not detected in WES data by ClinCNV.

| Category | Number of FNs |
| --- | --- |
| (Likely) artefact | 15 |
| (Likely) TP, but <50Kbps | 7 |
| TP (centromeric region) | 2 |
| TP (only one exon affected) | 3 |
| TP (6/60/131 exons) | 3 |

Finally, we have tested how high is ClinCNV’s Sensitivity in comparison with arrays when including off-target reads.

Our results showed that ClinCNV analysis of WGS data could replace platform-specific software array-based CNV detection in diagnostics. ClinCNV WES analysis without off-target reads misses all purely intronic/intergenic CNVs, with off-target reads still missing 15% CNVs (also mainly intronic or intergenic). The number of false negatives reduces to 5% for CNVs larger than 200k. We observed that arrays also miss some CNVs longer than 50Kbps (at least 3 cases within our test cohort) and are highly unreliable for CNVs shorter than 50kbp, producing many artefacts. We suggest that only whole-genome sequencing results should be used as the gold standard in future studies.

#### ii.3 Sensitivity of ClinCNV for detection of CNVs longer than 15kbp in shallow WGS data

Correlation of the number of detected CNVs and coverage depth is shown in Figure fig. 3.

**Figure 3:**
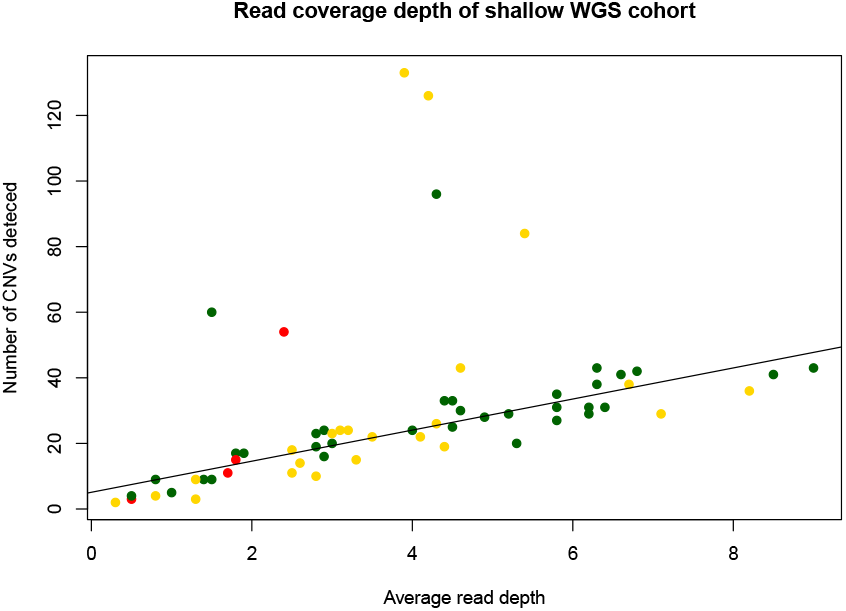
Analysis of Shallow WGS cohort: each dot is one sample, number of detected CNVs on y-axis, read depth on x-axis. Green color: CNV succesfully found. Yellow: CNV was not found, but its length was smaller than the detection limit (15Kbps). Red: CNV was detected as longer tha 15Kbps and not found by ClinCNV. Robust linear model shown to indicate the trend.

Four False Negative samples contained three or more CNVs or a very long CNV suitable for detection. However, neither a single one was detected by ClinCNV, nor any signs of decreased/increased coverage were observed. All of these four samples were analyzed with an external read-depth based CNV detection tool using Agilent SureSelect PathWay BRCA v2 panel (three samples) or ssHAE v6 (one sample), thus, it is highly likely that all the detected CNVs were False Positive in the primary analysis due to low Specificity of CNV detection in targeted sequencing.

All the CNVs detected by ClinCNV had concordant genotypes with CNVs detected by alternative methods (1 homozygous and15 heterozygous deletions, one homozygous and six heterozygous duplications).

We have shown that ClinCNV can be used to detect CNVs in shallow WGS samples, but the size of the detected CNVs is dependent on the actual depth of sequencing. For example, we identified three events of length ∼15 kbps in samples with coverage depths 6.4x, 6.3x, 4.5x. One CNV of approximately 30Kbps size was detected in the sample with 1.5x coverage. Since parameters of calling were not specifically calibrated for desired Precision / Recall, we manually checked all samples with CNVs shorter than 15Kbps in the genome browser, but no visually recognizable CNVs were identified.

#### ii.4 Expected number of CNVs in clinical settings

We describe the expected number of CNVs after “default” megSAP filtering for each type of analysis (WES, WGS, shallow WGS) as mode, minimum and maximum in the table 9.

**Table 8:** Comparison of array-based CNV calls with ClinCNV WES results (using both on-and off-target reads), 406 CNVs discovered with microarray technology were used.

|  | all | default |
| --- | --- | --- |
| % detected | 79.64 | 69.57 |
| % found, but CN mismatch | 4.33 | 3.05 |
| % missed | 16.03 | 27.48 |

**Table 9:** Visually defined minimum, mode and maximum of the distribution of number of highquality CNVs, per platform.

|  | Min | Mode | Max |
| --- | --- | --- | --- |
| WES | 43 | 87 | 141 |
| WGS | 578 | 950 | 1651 |
| shallow WGS | 47 | 148 | 286 |

### iii. Detection of CNVs in WES data

#### iii.1 Comparison between ClinCNV and ExomeDepth using research WES samples

Using the default parameters, the following results were obtained (shown in table 10).

**Table 10:** FDR of ExomeDepth and ClinCNV (raw calls), CC = ClinCNV, ED = ExomeDepth

|  | FDR, sites | FDR, CNVs | # sites |
| --- | --- | --- | --- |
| CC del | 0.559 | 0.094 | 900 |
| ED del | 0.149 | 0.005 | 130 |
| CC dup | 0.409 | 0.064 | 354 |
| ED dup | 0.067 | 0.001 | 182 |

As a first conclusion, we noticed that using ClinCNV on 40 relatively low-covered WES samples did not allow accurate statistical parameter estimation. When we used 155 samples from the same study including these 40 (results provided in Supplementary), the raw CNV site FDR was equal to 44.5% for deletion and 34.9% for duplication sites, using the same parameters. Thus, an increase in sample size leads to considerably better results for WES cohorts.

By default, the number of calls and FDRs of callsets obtained with ClinCNV and ExomeDepth were drastically different. In order to make them comparable, we had to select quality filtering parameters. For deletions, when we increase the Log-Likelihood score to 30 and the Log-Likelihood score per 1kb to 3, we get 0.143 FDR of sites but detect 238 deletions instead of 130 detected by ExomeDepth.

Increasing the threshold to 34, we detect a similar number of deletions but with even better FDR. Introducing of Log-Likelihood per kb metric is not artificial; it helps to resolve long variants with a small number of targeted probes in between the borders, which are almost always artifacts, a similar feature was implemented in ExomeDepth since version 1.05. However, it was not possible to establish a comparable threshold for ClinCNV to achieve a similar FDR for duplications found by ExomeDepth. We used a threshold of 23 for the selection of 179 duplication sites and still got worse FDR (table 11).

**Table 11:**
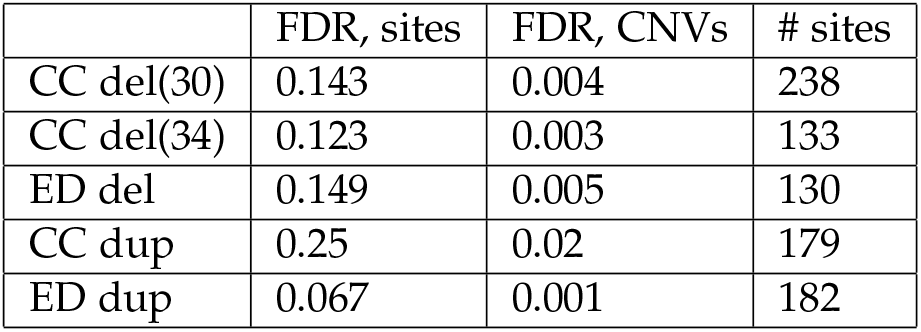
FDR of ExomeDepth and ClinCNV (filtered set), CC = ClinCNV, ED = ExomeDepth

**Table 12:** Comparison of ExomeDepth and ClinCNV using in-house samples, singleton variants.

|  | # sites | FDR |
| --- | --- | --- |
| ClinCNV del | 604 | 0.403 |
| ExomeDepth del | 473 | 0.428 |
| ClinCNV dup | 832 | 0.385 |
| ExomeDepth dup | 399 | 0.385 |

We checked how our filtered findings from two callsets with similar properties intersect. We used a relaxed threshold: two detected CNV sites from different callsets intersect if their intersection is bigger than the max(100 base pairs, 50% of the smallest variant). 56 of ExompeDepth deletion sites were mapped to 51 ClinCNV results, and 70 ExomeDepth duplication sites were mapped to 76 ClinCNV’s.

In summary, ClinCNV detects deletions in relatively low covered WES samples better, while ExomeDepth detects duplications better. We described the potential reasons for such discrepancy in Supplementary.

#### iii.2 Comparison between ClinCNV and ExomeDepth using well covered clinical samples

The results of calling for singleton CNVs are shown in table 12.

For all CNVs, the results were similar to the previous evaluation using research, relatively low-covered WES samples – ClinCNV detected the same number of CNV sites as ExomeDepth, having better FDR (1.316/1.315 deletions with FDRs of 0.364 and 0.39 and 1.049/1.002 duplications with FDRs of 0.313 and 0.377). From here, we can conclude that ClinCNV outperforms ExomeDepth., this time for both deletions and duplications. It may happen because, for high coverage samples and large cohorts, normal distribution approximation is accurate enough, while for relatively low coverage WES robust negative binomial model used by ExomeDepth is preferable.

#### iii.3 Detection of CNVs in clinical WES data in trios

The main interest in trio calling is the number of real *de novo* CNVs and the number of Mendelian errors. The plot in fig. 5 shows density of numbers of *de novo* CNV calls per sample (one exon or longer). Only 192 trios had less than 20 *de novo* CNVs with a quality score of 20 or higher (and thus formed this density). 43 trios with more than 20 *de novo* CNVs were excluded. The number of variants is decreasing rapidly with increasing the quality threshold. We can conclude that approximately two-thirds of the analyzed trios have less than 20 candidate CNVs even before diagnostic gene list filtering, which is realistic for further analysis, considering observed phenotype and other clinical data. For one-third of samples, potentially, higher quality thresholds can be applied. For samples with many CNVs detected, the resequencing of failed samples can be considered an option.

**Figure 4:**
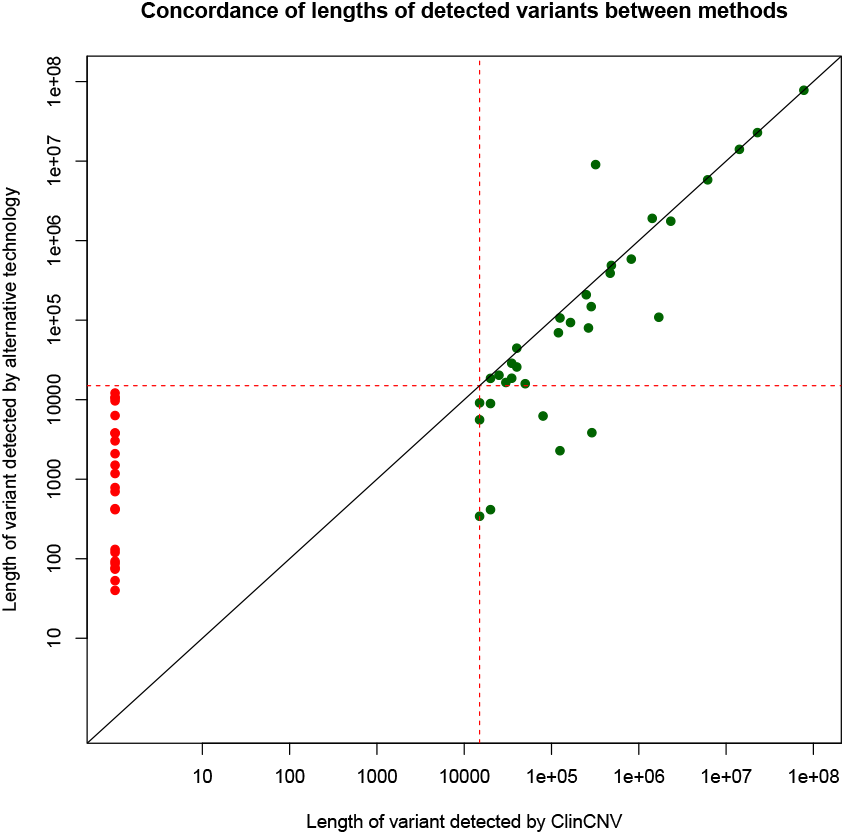
Overview of variants and their lengths detected and missed by shallow WGS analysis by ClinCNV. Red dashed lines denote detection limit, each dots is a CNV. Many CNVs which were previously detected as shorter than 15Kbps according to MLPA analysis (green dots below red horizontal line) were detected by ClinCNV since their true size was bigger.

**Figure 5:**
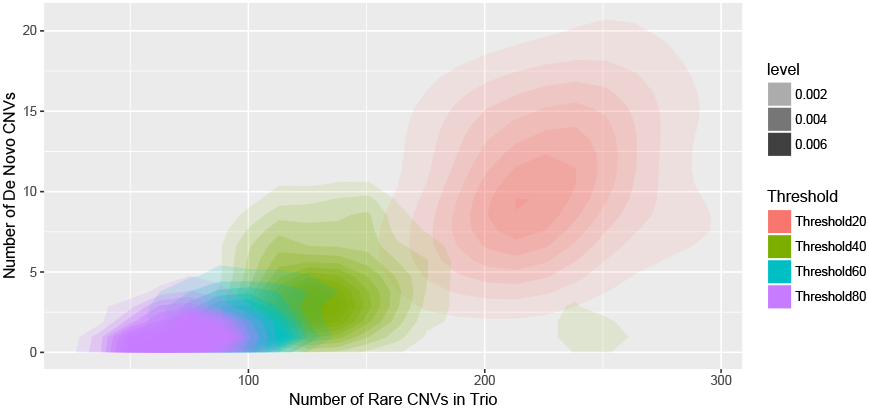
Density of number of CNV sites per trio vs amount of de novo CNV calls in a child.

IIt is well known that WES samples have a lot of technical artefacts, such as coverage outliers, which may be mistakenly detected as CNVs. Such extremely low or high coverage events 9 can not be filtered via increasing quality thresholds since they are extreme. We decided to repeat the analysis, counting only copy-number changes such as heterozygous deletion or duplication in regions normally diploid in parents since such events are more likely to be real. Comparing the new plot (fig. 6) with the previous one, we may conclude that around 5 CNVs per sample are highly likely to be technical artefacts (detected as homozygous deletions/duplications or higher copy-numbers). It further reduces the number of variants for clinical evaluation.

**Figure 6:**
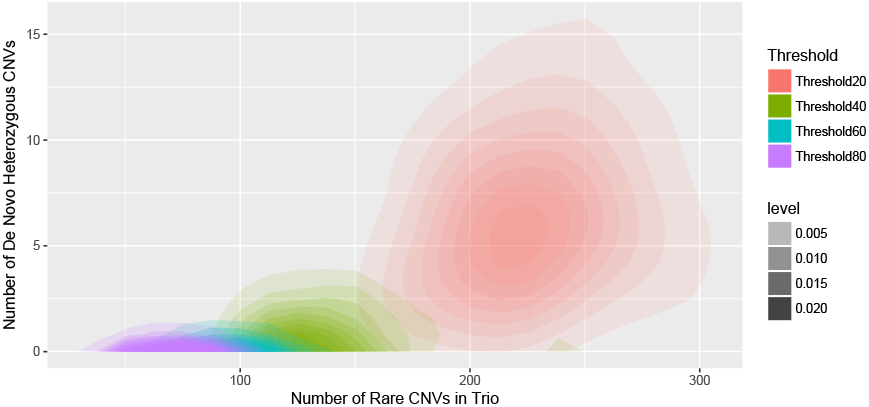
Density of number of CNV sites per trio vs amount of de novo heterozygous deletion and duplication calls in child.

In order to estimate the advantage of joint trio calling compared to single sample calling and merging of results, we estimated the density of the number of CNVs detected only in the child but not in parents (thus, it would be falsely considered as a *de novo* event). We can see that using a single sample calling, we detect fewer CNVs using the same thresholds (fig. 7).

**Figure 7:**
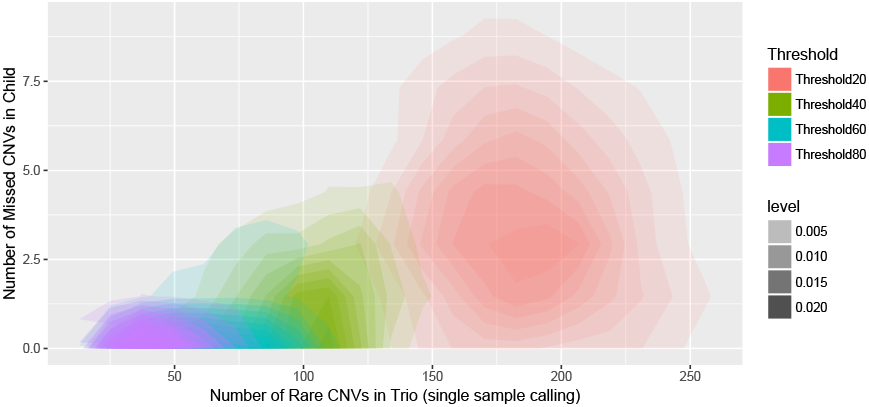
Density of number of CNV calls found in both child and one of the parents in joint analysis, but only in child in single sample analysis vs number of CNV sites detected using single sample calling.

Errors can be of another type – CNV is not detected in a child in single sample calling but presented in both child and one of the parents and recognised in joint trio calling. The average number of such errors is represented in fig. 8. Such errors may be clinically relevant, especially in the case of recessive diseases.

**Figure 8:**
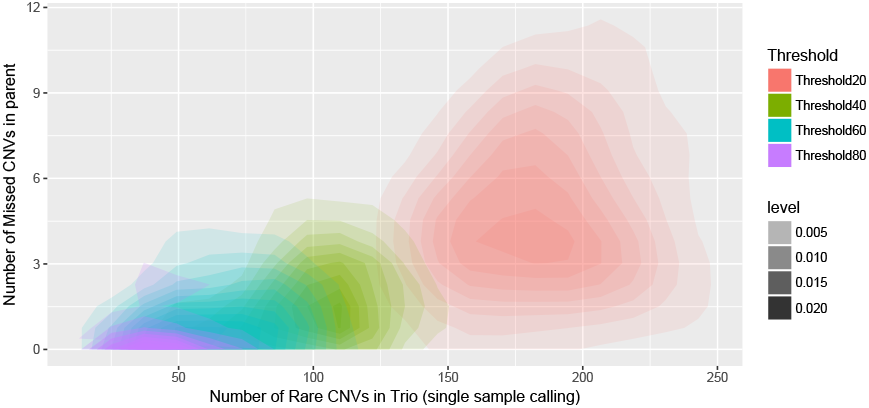
Density of number of CNV calls found in one of the parents and child according to joint analysis, but only in parent using single sample calling vs number of CNV sites detected using single sample calling.

QC failed (having too many CNV calls) trios must be analyzed carefully, using more strict thresholds. A QC failure of one out of three samples is enough to ruin the whole trio analysis. Accurate analysis is impossible for the case when the child sample has too many CNV calls, but for the parent with too many CNV calls, it is feasible unless the number of CNVs in a parent sample is of an order of thousandsin this case, real CNVs in the child can be “masked” by false-positive variants in a parent.

### iv. Estimation of quality of WES variants using array data and Quantile Random Forest Regression

As described in the introduction, a standard evaluation of CNVs called in NGS data with array data, namely, calling CNVs in microarray data and intersecting with NGS results, is sub-optimal. Assigning True/False positive labels for CNV calls is impossible for regions containing small numbers of markers or noisy regions. Thus, a more sophisticated method for performance evaluation was required.

The results (predicted and actual Precision and Recall) are shown at fig. 9 (internal validation of ssHAEv6 samples) and and fig. 10 (training in ssHAEv6, test with ssHAEv7 samples). Since no real True/False Positive labels were available and we always worked with probabilities, **the Precision-Recall curve estimation is not monotonic**. As we can see, the predicted (rainbow curve) and observed curves (black) match very well, which means the method we used was adequate to evaluate the Precision/Recall of each separate variant.

**Figure 9:**
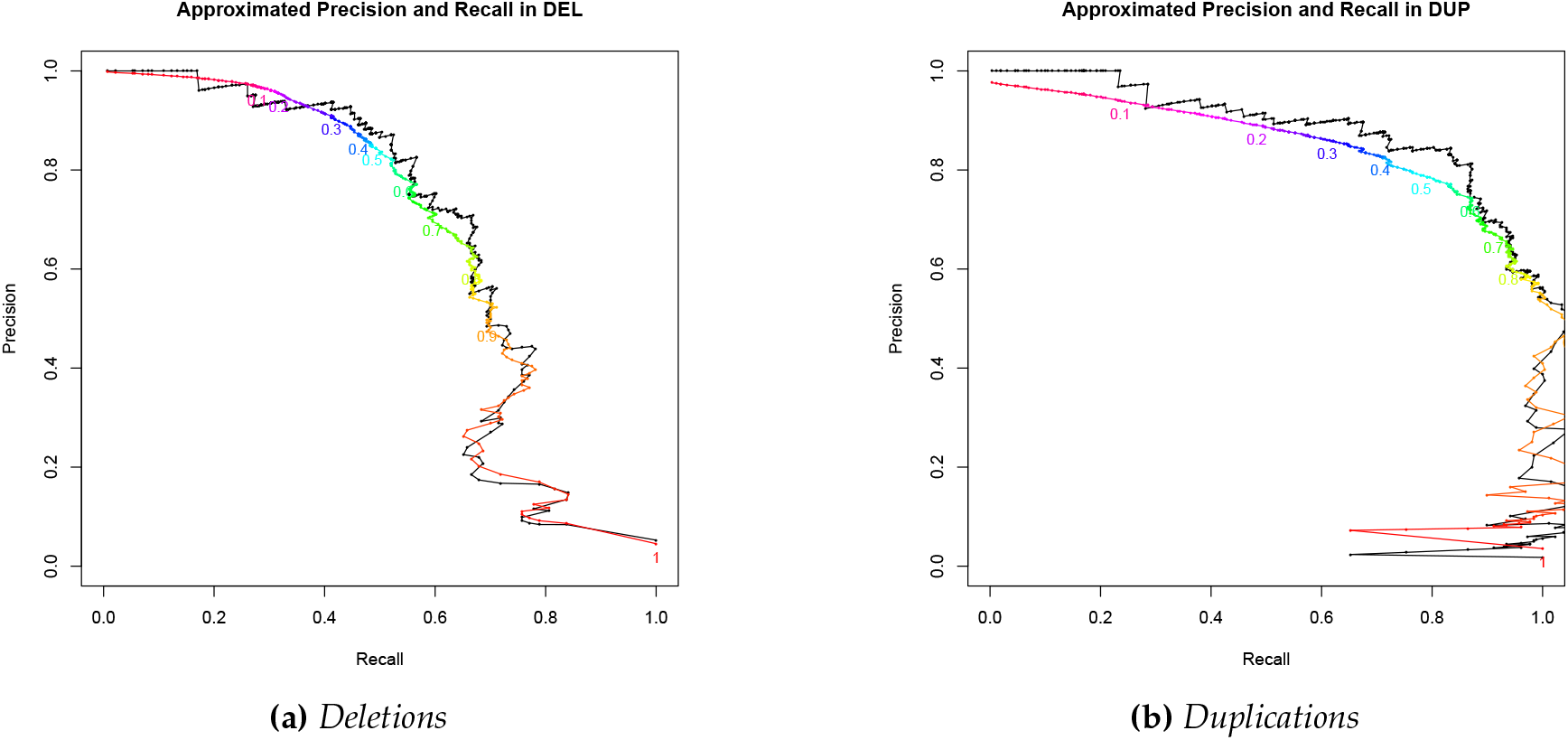
Estimated precision and recall for internal training and validation within ssHAEv6 cohort (60/40% split), black line - observed, rainbow line - predicted using machine learning technique. Numbers from 0.1 to 1.0 are showing the quality metric of each CNV, included into the cohort. Each dot denotes 0.002 step of allolwed individual CNVs’ FDR.

**Figure 10:**
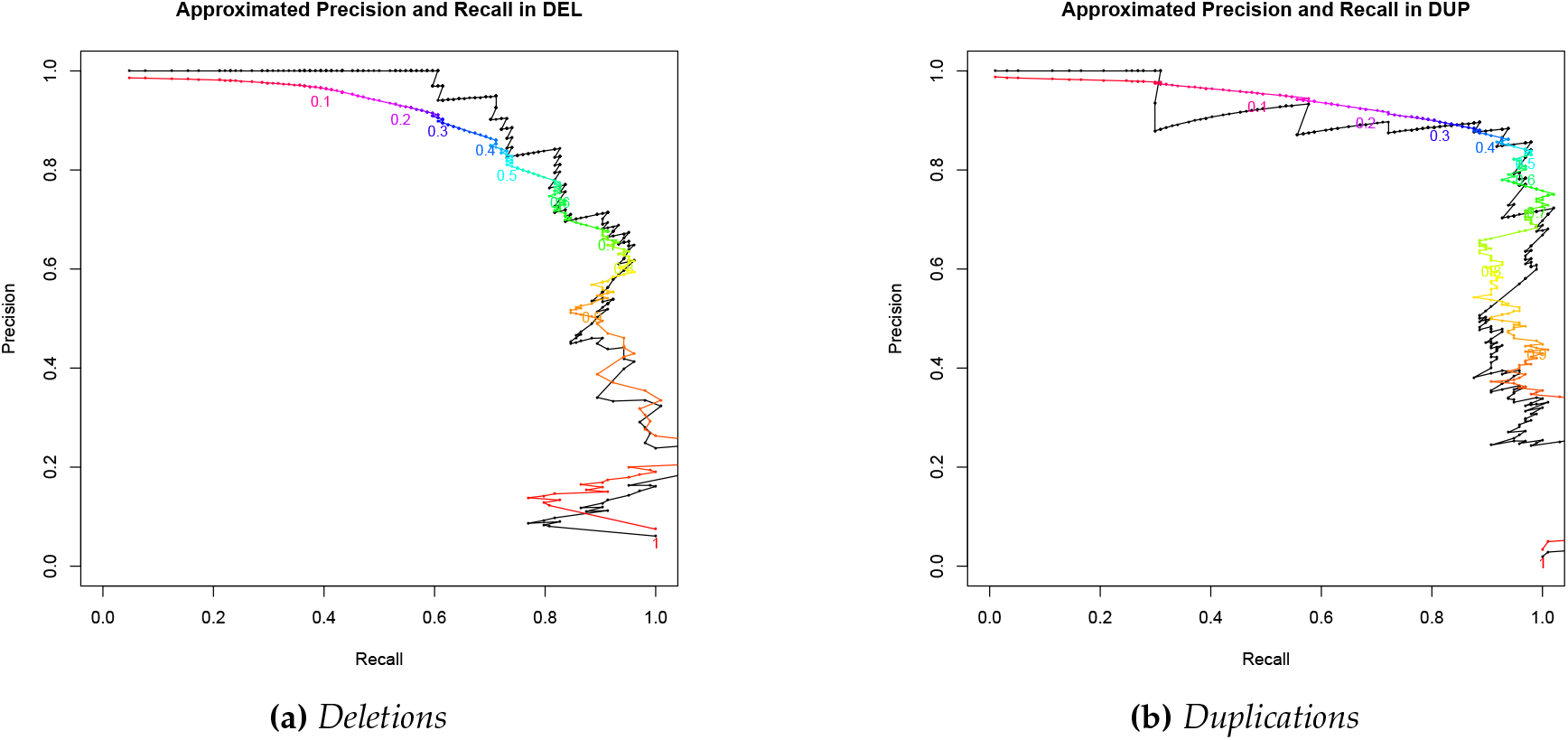
Estimated precision and recall for training within ssHAEv6 cohort and validation using ssHAEv7 cohort, black line - observed, rainbow line - predicted using machine learning technique. Numbers from 0.1 to 1.0 are showing the quality metric of each CNV, included into the cohort. Each dot denotes 0.002 step of allolwed individual CNVs’ FDR.

The estimation of Recall, unlike FDR, is more tricky since the distribution of p-values is not exactly uniform. It is especially true for the values in the bottom right corner (high Recall and low Precision) – due to the random chance alone, we could have higher or lower proportion of False Positive variants (p-value bigger than 0.5) than in the whole cohort, that is why the plots are largely unstable in this part and Recall may occur to be larger than 1. This volatility may be resolved via randomization and averaging. However, we show the original plots with one iteration of training and prediction. In general, these Recall values can be used as a rough approximation in order to understand where the curve “breaks”. It can not be used as a measure of the actual performance. Moreover, we tested the Recall of the CNVs that we could retrieve using ClinCNV with the relaxed threshold and, as described above, around 15% of array-detected CNVs that affect at least one exonic region were missed in the primary analysis.

Several differences between fig. 9 and fig. 10 worth to be discussed. At first, the predicted curves for deletions almost perfectly follow the curve of observed curves, but the Recall is smaller in internal validation within ssHAEv6 enriched cohort. It may happen due to 1) increased by more than 50% of training set size,2)increased quality of data for ssHAEv7 cohort due to both higher depth of sequencing and improved panel design. Also, our FDR seems to be overly pessimistic for deletions from ssHAEv7 cohort, which may be explained by the increased quality of calls or slight changes in target design (more probes are located in the tested regions). For the duplications predicted curve seems to be over-optimistic for ssHAEv7 samples. The large drops in Precision are likely due to the discreteness of our data and several unfortunate outliers since we have tried to analyse random sub-samples of ssHAEv7 cohort and, if averaged, there is no large drop in Precision as observed in the plot.

## IV. Discussion

In this paper, we have described the CNV detection pipeline used for calling from NGS data in clinical settings. The main conclusions are:

- paired-end mapping based and read-depth based methods should be used jointly for detection of CNVs in WGS data since the results they provide are not identical, even after controlling for FDR;
- microarray-based CNV calling, using detection of 50kbps CNVs as a criterion (genome-wide), can be replaced with high-coverage or shallow WGS-based, but not with WES-based analysis;
- the novel tool for read-depth based detection of CNVs in WES data does not outperform the existing well-established tool ExomeDepth in low coverage WES samples, but shows a similar performance. ClinCNV performs better than ExomeDepth in clinical-grade WES sequencing data;
- detection of CNVs in trios should be performed using either method that takes inheritance patterns into account (implemented in ClinCNV) or be performed with the relaxed thresholds per single sample and then merged to avoid false-negative results;
- calling of CNVs in clinical settings should not be concentrated on the False Discovery Rate of the whole callset, as commonly done in research projects. However, FDR for each variant and the additional considerations such as biological relevance should be included in the evaluation.

Our paper does not address the complex cases, such as complex SVs detection, mosaic CNVs detection, detection of variants in paralogous genes such as SMN1/2 or inside low mappability regions and comparison of the accuracy of calling between NGS-based and longread based methods. However, we provide a comprehensive evaluation of the developed tool in a wide variety of clinically relevant scenarios. Additionally, we provide guidance on the practical usage of our tool, quality control parameters and filtering.

## V Acknowledgements

Authors are thankful to Francesc Muyas for the fruitful discussions and to diagnostic team of IMGAG, UKT, especially Tobias Haack and Karin Schaeferhoff.

