## Supplementary Materials and Methods for "ClinCNV: multi-sample germline CNV detection in NGS data"

Stephan Ossowski[1,2]

[1] Institute of Medical Genetics and Applied Genomics, University of Tuebingen, Tuebingen, Germany
[2]Center for Genomic Regulation, The Barcelona Institute of Science and Technology, Barcelona, Spain
[3]Universitat Pompeu Fabra (UPF), Barcelona, Spain

June 10, 2022

#### I. Supplementary Methods

##### i. ClinCNV input data normalization

We decided to develop a data normalization procedure, aiming for the following goals:

1. We want to reduce the level of noise in data as much as possible without introducing unnecessary complexity since sophisticated methods for normalization may be computationally expensive, and it may become a limiting factor for the usage of our method.

2. Estimation of parameters has to be statistically robust. In other words – tolerate as many outlying values as possible since the read depth data is predisposed to have many artefacts even after carefully performed library preparation and sequencing. As a consequence, we had to avoid complex statistical models with the estimation of many parameters since it is technically challenging to control the robustness of such methods.

3. Normalized signals should not only tell if coverage was increased or decreased but also indicate the exact copy number. In essence, the property that the read depth in one copy of a particular region should be twice smaller than the number of reads sequenced from 2 copies should be preserved. Thus, methods such as Singular Value Decomposition (SVD) were inapplicable due to the fact that they "rotate" space of basis vectors, and distances there become hardly interpretable.

4. Usually, most of the genetic material remains copy-number state equal to the ploidy state (e.g., normally nearly all the genomic material in human autosomes is diploid), so it is not possible to directly estimate parameters for statistical models for each copy-number due to lack of data points for estimation. Having property 3), we can estimate mean levels of normalized read depth for different copy-number statuses having only data from

*Corresponding author

samples that have the normal number of copies, but we also need to estimate the variance in coverage depth for all the different copy-number models. We assume that read depth follows the Poisson distribution and uses square root transformation where possible since it stabilizes the variance [Anscombe et al., 1948].

5. During recent decades, the cost of sequencing dropped significantly, and it is much more common to have large cohorts of samples sequenced. Thus, we assume that the number of samples used for normalization is large. However, an important consideration is that even if a large cohort of samples were sequenced within the same sequencing facility and using the same procedure, it might happen that samples will still be affected by different batch effects. Therefore, one of the goals of normalization is to determine the most similar samples at first and then perform normalization within the cohorts stratified according to the sources of potential bias (typically, these sources are not observed).

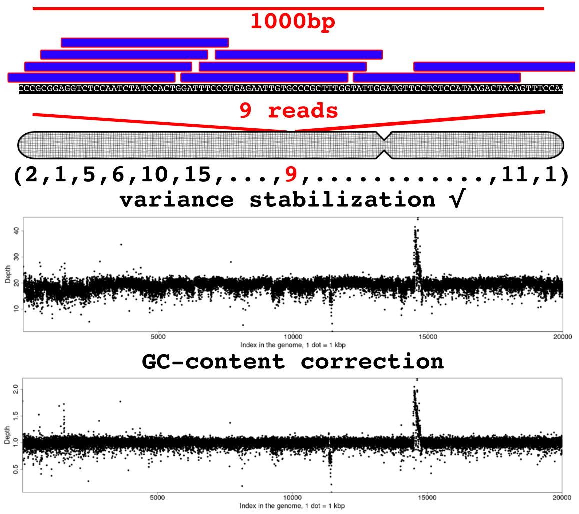


**Figure 1:** *Main steps of within-sample normalization used in ClinCNV.*

Using a restricted set of transformations for data normalization provides an opportunity to infer the expected amount of coverage increase. Variance stabilization transformation helps to infer variance in coverage without estimation of variance for statistical models of all copy numbers – we may estimate variance for the “majority of the samples” first and then assume that the coverage generated from different copies of the segment will have the same variance.

Homozygous deletions should be treated differently from other copy-number states in statistical modelling. The variability of the number of reads in homozygously deleted regions is not driven by the underlying amount of DNA, but the mapping process, which leads to non-zero read coverage even for missing parts of the genome. However, the inference of the statistical model for coverage in homozygously deleted regions is not straightforward, so we typically assume the mean level of expected coverage equal to 0 and variance equal to the variance of normalized coverage estimated for other copy-number states. We exclude data points that are closer to 0 than to the level of coverage expected for one-copy (heterozygous) deletions when we perform copy-number polymorphisms calling.

ClinCNV utilizes coverage depth values measured within windows (tiles) from the pre-specified set of intervals, which may be regions of enrichment and off-target regions for targeted sequencing such as TPS or WES or approximately uniform binning across the whole reference genome for WGS. Reads with low mappability (5 or less) usually are not counted (however, they have to be included in case the detection of variants within repetitive regions is preferred over the low level of stochastic noise). To correct for GC bias, we divide GC content into bins of similar GC content and use medians of coverages in such bins for normalization. Tolerance of GC content equality depends on the number of regions - by default, we round all the GC content up to 2 significant digits, but if less than 95% of the regions have at least 50 data points per GC value, we relax the tolerance value. The first steps of normalization are depicted on fig. 1.

We also noticed that for TPS and WES the lengths of the targeted regions are also a source of additional biases due to an uneven density of enrichment probes required to cover regions of different lengths. For example, using probes of uniform length X, we may expect all the collapsed (which means obtained as the overlap between all the targets in a particular part of the genome) regions with length less than or equal to X covered with just one probe, but for regions of length slightly more than X and less than 2X at least two probes are required which leads to increase of coverage in such regions. We correct such bias prior to GC content correction using locally estimated scatterplot smoothing (LOESS) using log2(length) as a predictor, which leads to up to 3% decrease in the individual sample's coverage standard deviation. This effect disappears for large regions or for coverage windows that are equally long across the genome. We filter out collapsed regions smaller than 50 bp since they often cause outlying coverage values.

Another strategy for target length induced bias in TPS and WES samples is binning collapsed regions into smaller overlapping windows. Due to the availability of enrichment probes and their intended targets for Agilent SureSelect Human All Exon enrichment kit (up to version 6) and several hundreds of non-tumor samples, we were able to investigate major sources of biases in sequencing results. As expected, we have observed peaks of coverage located at the centres of enrichment probes. We were also able to infer a strategy applied to the design of whole-exome enrichment kits produced by Agilent, and we have found that enrichment probes are designed in such a way that the difference between probes is equal to 80 base pairs while the length of probes is equal to 120 base pairs. A similar discovery was made by [Parrish et al., 2017] where the performance of ExomeDepth tool [Plagnol et al., 2012] was the best using the coverages summarized in windows of the size of 120 base pairs which is equal to the used enrichment probes' length. We have tried to divide targeted enrichment regions accordingly and analyzed several hundreds of samples. For each consecutive pair of windows we divided read coverage into three parts: coverage from non-overlapping part of the left window, coverage from the overlapping part and coverage from the non-overlapping part of the right window. We calculated coverages for non-overlapping parts and divided the coverage from the overlapping parts proportionally to the coverage of non-overlapping windows' parts. We concluded that the normalizations such as GC-content normalization worked more efficiently after such binning and larger parts of the enriched regions became available for the analysis with ClinCNV (since we filter out all windows with less than 50 regions with exactly the same GC). However, the results of subsequent CNV calling and the manual examination of detected variants that are located within the borders of the collapsed target regions showed that most of the CNVs detected were not supported by other evidence such as the presence of split-reads or wrong paired reads orientation. The absence of such alternative evidence, as well as the unexpectedly high number of homozygous deletions detected, indicated that the vast majority of findings are most likely false positives caused by technical issues of library preparation and sequencing. Thus, we were not able to conclude that such a strategy improves the results sufficiently. Since the binning procedure creates an additional step in the bioinformatics pipeline, which is quite a time consuming, we used the previously described strategy for length induced bias correction.

In order to be able to work with sex chromosomes, we infer the sampled individual's gender during the first steps of our algorithm, calculating medians of X and Y chromosomes' coverage and running k-means with two expected components centred at (1,0) and (0.5, 0.5) and assign sex as "female" or "male" for samples from these 2 clusters, respectively.

##### i.1 Removing batch effects via clustering

Batch effects in read coverage depth data may be removed using several statistical techniques, such as SVD [Krumm et al., 2012] or PCA [Fromer et al., 2012], but it is relatively difficult to control the robustness of such methods in different types of data. In particular, a few short CNVs or a small number of bad quality samples can not affect most of the batch effect correction techniques, but if a large fraction of a genome is altered by CNVs due to the presence of aneuploidies, many short CNVs or a significant amount of noisy samples – all such factors may have a dramatic effect on batch effect removal efficiency. Instead, we propose a clustering method for the separation of sub-groups of samples with similar patterns of technical variation and performing analysis within groups of similar samples.

For coverage normalization for CNV calling in parent-child trios, the desired property is to analyze all three samples within the same cluster. In the vast majority of cases, this requirement is automatically satisfied since trios are usually prepared and sequenced at the same time; however, in practice, there were some exceptions. To correct the clustering of such samples, i.e., to force all 3 related samples from the trio to end up in the same cluster, we choose median coverage value for each genomic region from these 3 samples, add small random noise and assign such coverage profiles to all the samples from the trio. This replacement of actual coverage values with the median-smoothed profile across the trio is used for clustering only.

An important parameter that has to be selected for each dataset separately is the minimum number of samples inside each cluster. Clustering into smaller groups of highly similar samples is better for batch effect removal, but the decrease in sample size increases errors in the statistical estimation of parameters. We would recommend choosing one-third of the total amount of samples and increase/decrease this value for more fine-tuning using plots produced by ClinCNV (fig. 2), as guidelines. For relatively small datasets (less than 60 samples), we would recommend to completely skip the clustering step.

Before clustering, we try to remove the potential impact of polymorphic regions. Since a large part of the human genome experienced copy-number changes in polymorphic regions, which are not representative for estimation of technical variability patterns, and low-variability regions are not informative for the estimations, we filter out all the regions with variability in top or bottom 20%. Then we smooth coverage profiles using the rolling median (the default length of the rolling median is five regions). Some samples may have a significant amount of outliers which are typically very short (1-3 consecutive regions), and their effect is minimized after the smoothing.

The matrix of distances between smoothed coverage profiles is calculated next. By default, Manhattan distance is used; however, for very large datasets (more than 1000 samples), correlation-based distance showed better and easier separation between samples of different coverage patterns.

Next, we perform isometric multidimensional scaling (MDS, [Venables et al., 2002]) mapping of smoothed coverage profiles. We use three components for mapping since it provided better results and faster convergence compared to two-component MDS. For the last step – clustering – we use simple hierarchical clustering (Ward method). We calculate the matrix of Euclidean distances between coordinates of MDS-mapped coverage profiles of samples and perform hierarchical clustering, increasing the number of clusters by one at each iteration, starting from 2. When less than 80% of samples are clustered into subgroups of pre-specified minimum size (which means that more than 20% of samples are clustered into clusters of a smaller size than the pre-specified threshold), we stop increasing the number of clusters. We keep only clusters bigger than the minimum size and assign samples from smaller clusters to the closest large sub-groups.

Summing up, 1) we cluster all the samples according to the similarity between their cover-

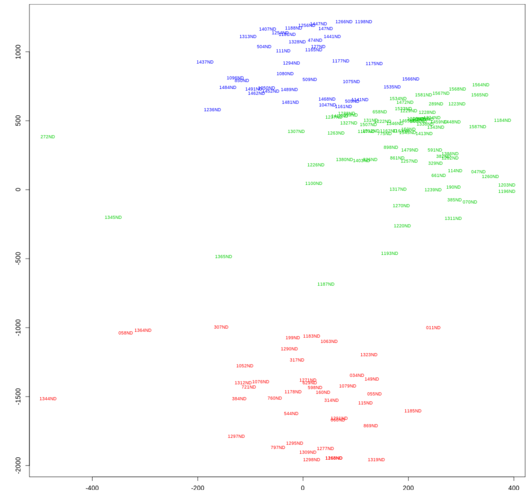

**(a)** *Clustering of germline WES samples from CLL study (results of CNVs calling in this cohort are described in the next part of the thesis). Outlying samples do not form separate small clusters.*

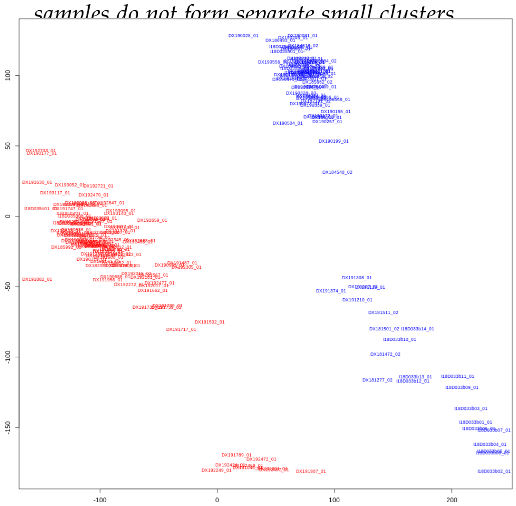

**(b)** *Clustering of panel sequenced samples. Visually we can identify 4 clusters, but amount of samples in 2 of them is small. In order to keep statistical parameters estimations accurate, we separate samples into 2 big clusters.*

**Figure 2:** *Examples of clustering of similar samples.*

age profiles, 2) all the output clusters are larger than a pre-specified size (i.e., outlying samples do not form separate small clusters), so we control sample size for statistical estimation of various parameters. We run normalization and calling within the groups inferred with this procedure separately.

#### ii. Graphical depiction of how algorithm works

The graphical example of how the algorithm finds one CNV is shown in fig. 3. The actual number of states is bigger (normally from copy-number 0 to copy-number 8) or, for common CNVs, the states may be "there is a common CNV site, modelled with normal mixture" or "there are just several outliers".

##### ii.1 Quality metrics, produced by ClinCNV

As an output quality value, ClinCNV provides the log-likelihood score for the variant. However, since we always perform multi-sample calling, it is possible to provide additional quality parameters, such as a number of samples that may also have CNV at this position (log-likelihood is bigger than 1, not to be confused with the actual allele frequency of a variant – may indicate both presence of CNVs and noisiness of the region), q-value (FDR-corrected p-value, obtained from t-test), log-likelihood per 1 KB and log-likelihood per data point (these are usually the same for WGS samples with coverage depths pre-calculated in 1KB windows, but they differ for WES or WGS with different window size), and others. These and other quality metrics are used for making a classification of variants into True Positives and False Positives using the Random Forest method.

### II. Supplementary Results

#### i. Common CNVs regions

The length of the CNV database, generated from a rare CNV detection algorithm and our in-house WGS cohort (280 samples), was

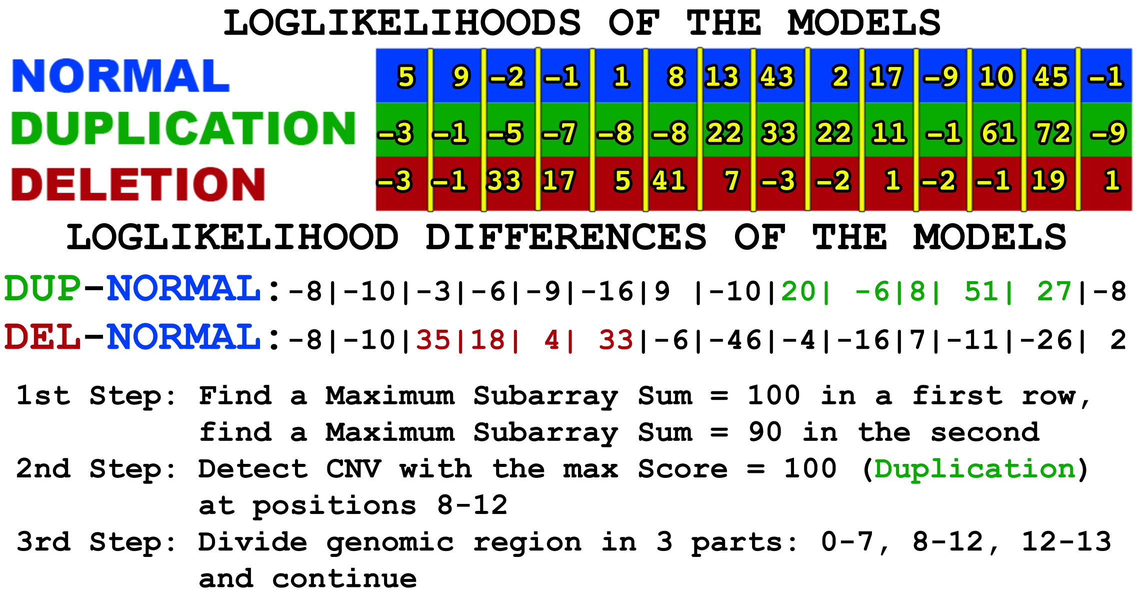


**Figure 3:** *Toy example on finding one CNV with the help of matrix of likelihoods.*

equal to 73.323kbps. The one detected with common CNVs detection algorithm, and a large cluster of similar samples from Pan-cancer analysis of whole genomes (PCAWG) cohort [Campbell et al., 2017] was equal to 97.256kbps. The intersection between common CNVs regions between 2 datasets was equal to 42.455kbps. Such a large difference may be explained by several reasons: 1) common CNVs detection algorithm has the power to detect some common CNVs which are smaller than 1kbps while rare CNVs detection algorithm is unlikely to find them due to large variance, 2) frequency of common CNVs in our cohort was set equal to 2% while for PCAWG cohort we had 2.5%, 3) due to random fluctuations and the fact that PCAWG cohort included people of different ancestries while our in-house cohort was recruited in Germany the actual content of common CNVs could change, 4) both datasets included some "false-positive" common CNVs due to the potential batch effects of sequencing and these batch effects could be different.

In general, for clinical diagnostics, we would recommend using the database of "common" CNVs obtained from our in-house samples. Ultimately, we are interested not in the fact that this CNV is common in the studied population but in how many times the algorithm detected it so we can exclude both common CNVs and artefacts of sequencing. Regions of common CNVs with high allele frequency (more than 75% of samples have copy-number different from 2) have to be analysed with a common CNV detection algorithm - they are almost undetectable by rare CNV detection methods.

However, if the goal of the research is to correlate common CNVs with the phenotype, we would recommend using the genotyping using the common CNVs detection algorithm. Even the data has to be prepared differently for these approaches: for the rare CNVs detection, one may want to keep reads with low mappability so they may indicate problems with paralogous genes, while for common CNVs detection, they have to be filtered out for accurate genotyping. A rare CNVs detection algorithm may tolerate the presence of batch effects in small parts of the samples, while common CNVs have to be detected in the cohort of highly similar samples.

#### ii. PCAWG supplementary results

##### ii.1 CNVs and separation of samples of different ancestries

A scientific question we wanted to answer is if the frequency of CNVs is different for some particular predisposition genes and some specific cancer types compared to other types of cancer, e.g., it is well known that BRCA1/2 deletion in the germline is a risk factor for breast adenocarcinoma development. To do so, we had to separate the populational effect on CNV frequencies from CNVs associated with cancer type. We had ancestry information for 1762 samples (they were classified as "AFR", "ASI", "EUR"), and 709 were marked either NA or "Others".

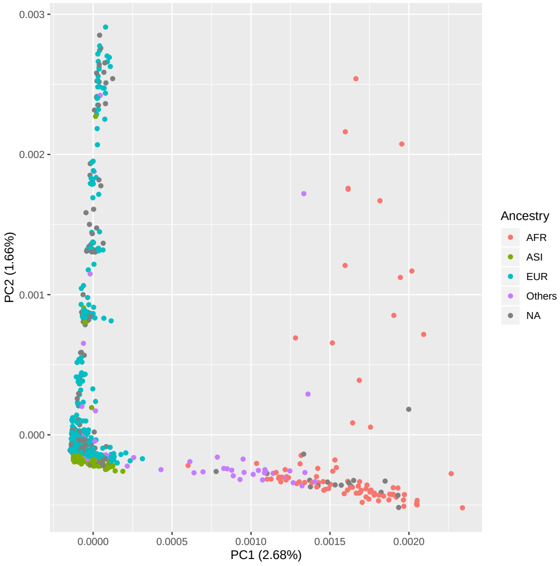


**Figure 4:** *First 2 principal components plot based on rare CNVs detected by ClinCNV.*

As can be seen from the PCA plot in fig. 4, even for rare CNVs population stratification exists, and since some cancers have been analyzed almost solely within one population, this factor had to be accounted for. We decided to impute the ancestry of these 709 samples even if we had only rare CNVs. We divided 1762 samples with ancestry information available into a train set (80%) and test set and trained a random forest classifier, using the presence or absence of CNVs as a predictor, and only CNVs that occurred twice or more per studied cohort were used. The classifier showed good accuracy on the test samples: only one sample out of 352 from the test cohort was misclassified (a sample with recorded Asian ancestry was classified as European ancestry sample). We have applied trained classifiers to our 709 samples of unknown ancestry and ended up with 126 samples recognized as African ancestry, 371 recognized as Asian ancestry and 1974 as European ancestry. We extracted the list of cancer predisposition genes from [Whitworth et al., 2018]. 56 genes out of 133 were altered in at least one sample in our cohort. We tested proportions using the Cochran-Mantel-Haenszel chi-square test for each cancer type separately, counting all other cancer types as a control cohort. However, none of the results remained significant after multiple test correction, even if many genes were significantly enriched when we used Fisher test without population-based stratification. Thus, we can conclude that despite the comparatively large number of samples, the power was not enough to detect any kind of enrichment of germline CNVs in specific cancer predisposition genes.

Analyzing such a diverse cohort with a huge shift in the number of patients of European ancestry, we realized that the "common CNV" term could not be applied to the pan-ancestry studies, especially imbalanced. E.g., these are the plots of lengths/numbers of CNVs per sample from different populations (fig. 5) and amount of genetic material varied was much higher in non-European ancestry samples since 1) human genome hg19 was based on, most likely, European genomes, 2) common CNVs that were excluded at the first step of ClinCNV algorithm were detected in a sub-sample with the vast majority of samples being of European ancestry. More sequencing projects involving non-European samples are required for a more accurate analysis of large genomic variants.

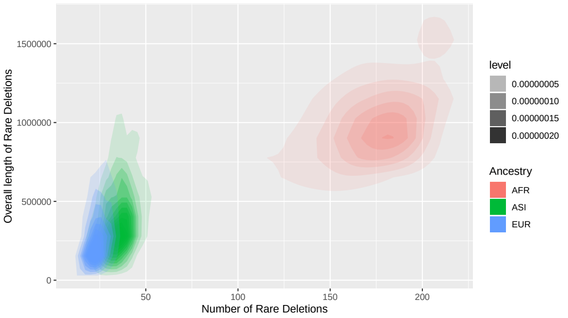


**(a)** *Lengths and number of deletions, by different ancestry.*

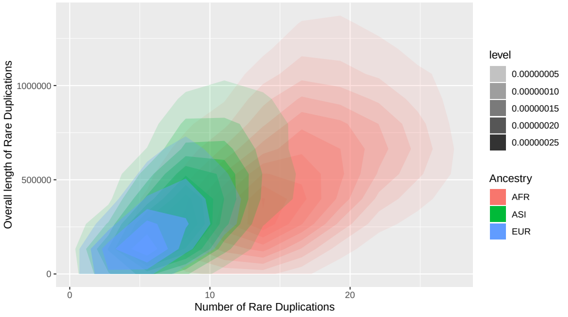


**(b)** *Lengths and number of deletions, by different ancestry.*

**Figure 5:** *Overview of CNVs numbers detected in samples of different ancestries.*

##### ii.2 CNVs detected by ClinCNV in cancer predisposition genes (PCAWG cohort)

Cancer predisposition genes, affected by CNVs, are presented in fig. 6. The largest number of variants was observed in the genes ALK, CDH1 and EGFR. BRCA1 was affected only 3 times and BRCA2 was not affected even once. DELLY detected only one deletion in BRCA1, which shows the usefulness of ClinCNV for CNV detection even if the alternative PEM method is applied.

##### ii.3 Comparison between NGS-based and array-based detection: additional plots

Lengths and numbers of variants for arrays and ClinCNV detected variants are provied in fig. 7 and fig. 8

##### ii.4 Results of CNV calling in whole-exome sequencing data of Chronic Lymphocytic Leukemia

We have tested CNV calling with ClinCNV in WES data from 435 samples from the ICGC Chronic Lymphocytic Leukemia (CLL) project. The cohort data, including clinical and genomic data, is fully described in [Puente et al., 2015]. Two panels were used for sequencing: Agilent Sure Select v4 (51MB) and v5 (71MB). The panels were not only different in size but also the median coverage was substantially different (fig. 9). Thus, we analyzed these datasets of sizes 274 and 161 separately. Interestingly, these two datasets produced different results from a technical point of view. The median coverage was lower than the coverage of WES samples sequenced nowadays for clinical purposes (80x-100x). However, these samples had array data for the validation, and thus, we decided to include them in benchmarking and analysis.

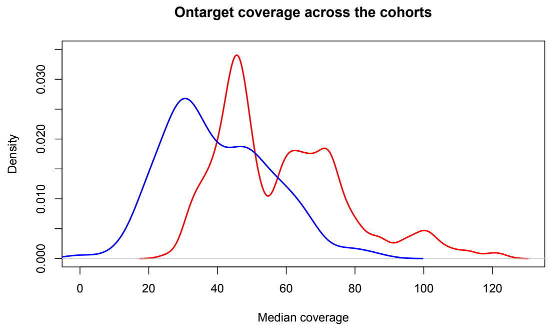


**Figure 9:** *Median on-target coverage of samples from v4 cohort (red) and v5 cohort (blue).*

For both datasets, the same rule was used for QC filtering. We plot a density for the number of CNVs and choose a threshold using a shoulder rule - if the number of CNVs is bell-shaped, but at some point a "tail" starts to form, we consider all samples with the number of CNVs in this tail as QC failed. We used 100 as a threshold for the number of CNVs for 161 samples sequenced with v5 and 220 as a threshold for the number of CNVs for 274 samples sequenced with v4 panel. So, the second

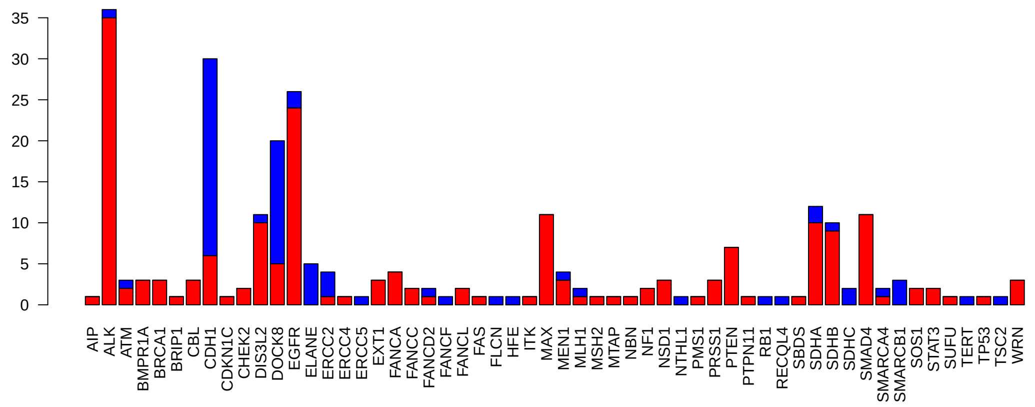


**Figure 6:** *Barplot of CNVs in cancer predisposition genes in the studied cohort. Red denotes deletions, blue - duplication.*

cohort, sequenced around 2x deeper, provided much more results, even if the panel was 20MB smaller. 155 and 265 samples remained for the germline calling. Variants with a q-value bigger than 0.05 were filtered out.

Almost all the samples had array data, so we performed the same intensity-based rank testing of the CNV callset as described in the previous section. As before, we have validated CNV sites, not CNV calls (however, there were many CNV sites that were represented by only one CNV call, so in this case, it is the same). Standard calling parameters were used (20 log-likelihood score as a threshold, 1 region as a minimum size of a CNV). "Sensitivity" is always a relative value in our analysis. It means a proportion of True Positive variants detected regarding the overall number of True Positive variants *detected by ClinCNV*. No gold standard CNV callset was available.

##### ii.5 Results of CNV calling in 265 Agilent SureSelect v4 Exomes

To check genotyping accuracy, we made a plot of array intensities of confirmed CNVs, according to ClinCNV's genotype (fig. 10). As can be seen, copy-number 0 is not always lower than copy-number 1, which highlights that homozygous deletion signature (absence of coverage) may be caused just by problems with hybridization. Copy-numbers 5, 6 and 7 have not had enough points for estimation.

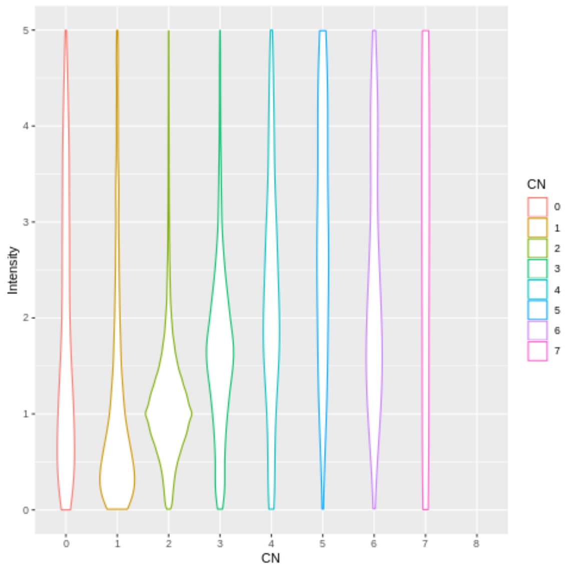


**Figure 10:** *ClinCNV's determined genotype vs array intensitiy for all variants with p-value less than 0.01.*

At first, we provide the raw estimation of FDR for different types of variants without any additional filtering. It is calculated as two multiplied by the number of variants suitable for evaluation, which had p-values bigger than 0.5, divided by the total number of variants suitable for evaluation. Two types of FDR may be assessed: FDR of the site and FDR of the variants. The second metric is calculated in the

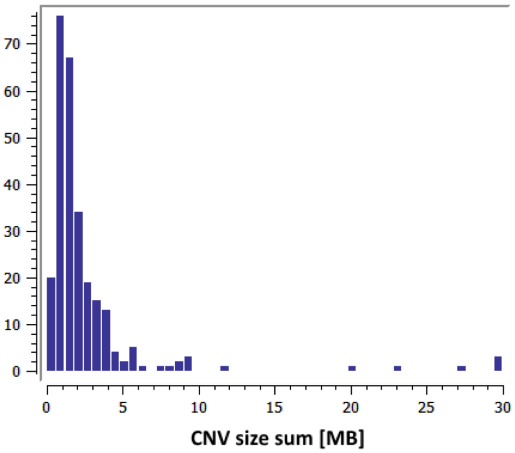


**(a)** *Length of CNVs per sample (arrays)*

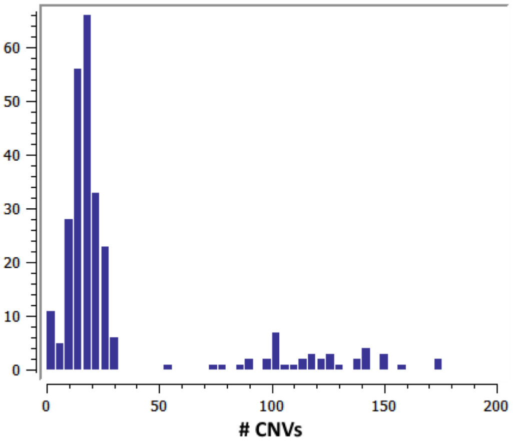


**(b)** *Number of CNVs per sample (arrays)*

**Figure 7:** *Number and lengths of CNVs per sample detected by array-based analysis*

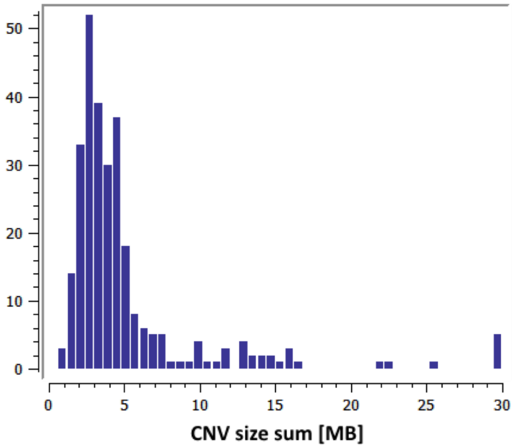


**(a)** *Length of CNVs per sample (ClinCNV)*

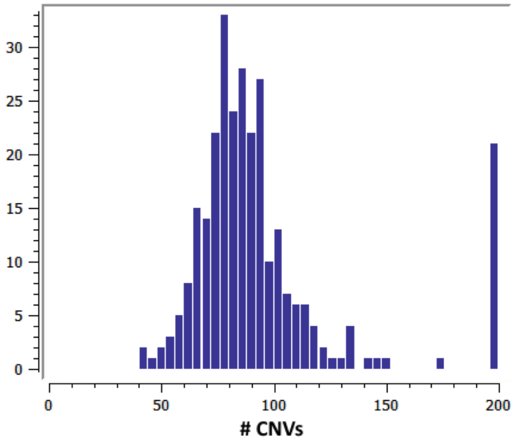


**(b)** *Number of CNVs per sample (ClinCNV)*

**Figure 8:** *Number and lengths of CNVs per sample detected by ClinCNV*

same way but takes into account the quantity of the variants that occur at the same site. The raw CNV site FDR was equal to 45.6% for deletions and 44.0% for duplications. The FDR of variants was estimated as 21.8% and 23.3% for deletions and duplications, respectively. For research purposes, FDR has to be decreased, and thus, these variants have to be filtered.

We have trained two random forest classifiers for deletions and duplications. Using 0.89 of random forest predicted probability as a threshold for a "True" CNV class gave us a Sensitivity of 0.591 and Specificity of 0.976 at the test set. For duplications 0.93 threshold was used, and Sensitivity of 0.574, Specificity of 0.968 were obtained. False Discovery Rate should be estimated twice as high as provided numbers (thus, one minus Specificity, multiplied by two, so FDR is expected to be around 0.05 for both types of CNVs).

418 duplication sites and 459 deletion sites were validated at approximately 0.05 FDR.

##### ii.6 Results of calling in 155 Agilent v5 sequenced samples

A plot of array intensities of confirmed CNVs, according to ClinCNV's genotype fig. 11. Inter-

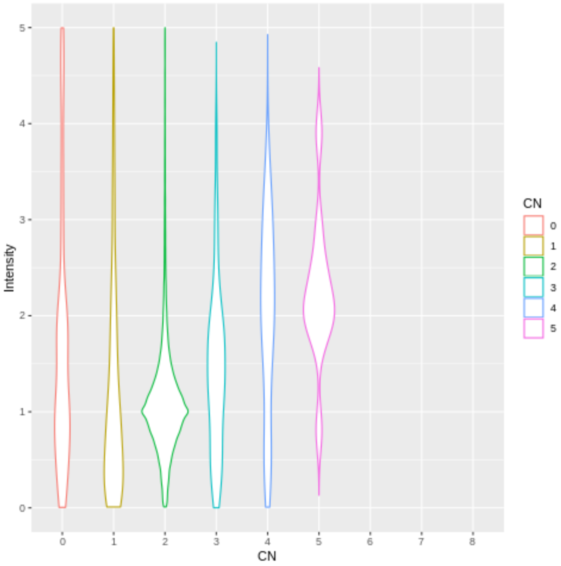


**Figure 11:** *ClinCNV's determined genotype vs array intensitiy for all variants with p-value less than 0.01.*

estingly, quite a lot of variants were called copy-number higher than 5. Nevertheless, none of them were validated with arrays. Again, we see that homozygously deleted regions do not necessarily correspond to lower array intensity. Thus, the absence of coverage in WES should be considered not perfect evidence of a homozygous deletion.

The raw CNV site FDR was equal to 44.5% for deletions and 34.9% for duplications. The FDR of variants was estimated as 21.3% and 23.5% for deletions and duplications, respectively.

We have trained two random forest classifiers for deletions and duplications. Using 0.92 of random forest predicted probability as a threshold for a "True" CNV class gave us Sensitivity of 0.76 and Specificity of 0.938 at the test set. For duplications, 0.6 was used as a threshold, and a Sensitivity of 0.76, Specificity of 0.94 were obtained, which gives a true false discovery rate of around 12% for duplications. Further increase of parameters did not provide us with a satisfactory FDR on the test set (Sensitivity drops too fast). The most probable reason for that is the small number of variants for validation, so the random forest was not able to extract true dependencies between variants' properties and the labels. Due to these reasons, we decided to check if the FDR would be stable if we selected another test/train sets, and the FDR was approximately at the same level.

401 duplication sites and 146 deletion sites were validated at the above mentioned FDR.

##### ii.7 Comparison of the callsets

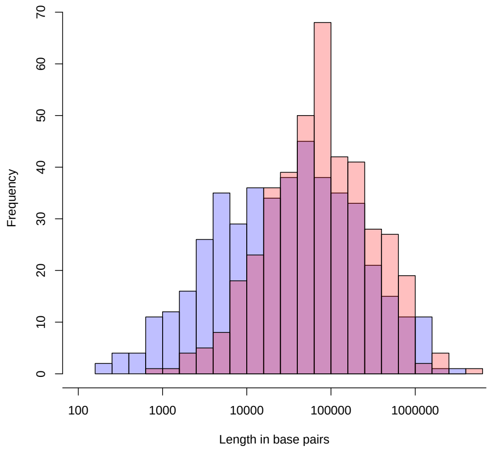


**(a)** *Length of CNV sites (v4, 5% FDR).*

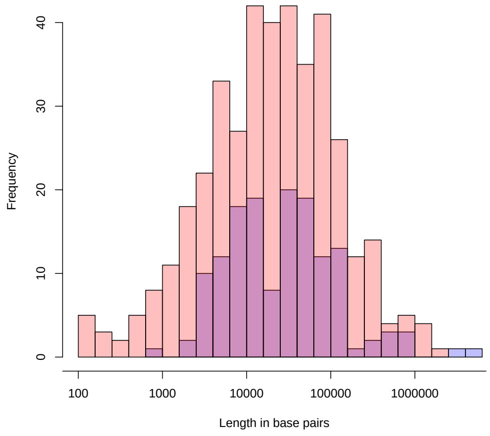


**(b)** *Length of CNV sites (v5, 12% FDR).*

**Figure 12:** *Overview of deletion and duplication site length in 2 WES callsets. Deletions are shown in red, duplications – in blue.*

Since these 2 callsets could not be directly merged, we provide the statistics separately. At first we show distribution of lengths at fig. 12.

Since we have not performed polymorphic regions filtering, we may expect many of the variants to be copy-number polymorphisms (occur more often than our pre-defined allele frequency of 2.5%). The plots in fig. 13 shows the frequencies of variants.fa

Analysis of recurrent CNVs in the germline of CLL patients (fig. 14), showed that ERCC2 was duplicated four times and one time in the independently analyzed datasets, as well as PMS2 (two and one duplications). Multiple deletions in PRSS1 (size plus three) were detected in both datasets.

#### iii. Difference in performance between ExomeDepth and ClinCNV

The ExomeDepth callset was even more strange since the author claims that he usually obtains around two-thirds of detected variants as deletions and we see the completely opposite situation in our dataset. However, no errors in the code were found in the ExomeDepth pipeline that we have used.

Such estimations of FDR may be inaccurate due to the small number of events tested – actually, we can not rely on the uniformity of p-values distribution, only 5/7 duplications/deletions were bigger than 0.5 in ExomeDepth dataset, and 11/5 was bigger than 0.5 in ClinCNV dataset, so differences in performance may occur by chance. Due to the large variability of FDR estimations, using this comparatively small callset, we decided to check the proportion of CNV sites that have p-values less than 0.05 in our callsets. For deletions, it was 0.81 for ExomeDepth and 0.77 for ClinCNV, and 0.79 and 0.66 for duplications, which is in line with the estimated FDRs.

The reference manual of ExomeDepth states that it may miss common variants and is more suitable for detecting variants that are presented in a single copy. We calculated the number of singletons detected by each tool. The filtered callset from ClinCNV was used. 128 deletions from ClinCNV were singletons, while only 68 of the ExomeDepth deletions were detected only once per cohort. The same number of duplication sites – 55 – was represented in more than 1 sample.

Surprisingly, many of the duplications, detected by ExomeDepth, showed no or very weak evidence in ClinCNV's coverage track in IGV, even very long ones (more than 50 exons). The very weak evidence means the segment is more likely to be normal (estimation closer to copy-number two than to higher copies). Yet, most of the checked duplication segments had elevated coverage (indicating around 2.1-2.5 "copies"). It could occur if the duplications were located in CNP regions which are not diploid in the population. We have checked and found out that 77 of ExomeDepth duplication sites were located in such regions (have more than 50% intersect of the variant's length), but 62 ClinCNV duplication sites were also located in such regions. Another explanation was that these variants are located in regions of low mappability, and reads with mapping quality below five were filtered as a preparatory step for coverage counting in ClinCNV, while ExomeDepth uses reads with mapping quality larger than 20 for the counting of reads. It may also explain the fact that ClinCNV detects more deletions at a similar FDR level. However, several large duplications detected by ExomeDepth were located in the regions of good mappability and still showed no evidence of duplication in particular samples visualized in IGV using ClinCNV's coverage track. Some duplications were located outside of the targeted regions provided by the manufacturer. ExomeDepth performs counting using the set of exons, not the targeted panel; thus, ExomeDepth has a signal in places where we did not calculate the coverage.

The last possible explanation for such discrepancy was in the nature of the data. Our data come from sorted blood cells, and many mosaic events may happen there. ClinCNV was created in a way, so it does not call mosaic variants if it is not explicitly asked to. We have compared the number of genes that are typically rearranged in blood cells between the callsets. ExomeDepth detected 18 CNV sites within HLA regions, ClinCNV detected only 6. 2/7/3 sites affecting IGK/IGH/IGL genes were detected by ClinCNV, 4/12/3 by ExomeDepth, respectively. Overall, ExomeDepth detected 37 variant sites in immunoglobulin and major histocompatibility complex regions, while ClinCNV detected 18. They were deletion and duplication sites in approximately equal proportion, so it cannot explain the large advantage of ExomeDepth in duplications calling.

A large amount of intersecting sites within one dataset may occur due to the over-segmentation and incorrect merging of the variants. ExomeDepth had 47 self-intersecting duplication sites within the dataset and 57 deletion sites. Our callset had ten self-overlapping deletion sites and 38 intersecting duplication sites. Thus, we can hypothesize that ClinCNV outperforms ExomeDepth in terms of breakpoint resolution, but the numbers are not big enough to explain the difference in calling fully.

It may be hypothesized that ExomeDepth outperforms ClinCNV due to the whole-sample beta-binomial model fitting, which may be more powerful. Using results of these 40 samples, but called with the whole cohort of 155 samples, does not confirm that, even if the results actually improve (table 1).

#### iv. Overview of platforms and variant length for shallow WGS analysis

##### iv.1 Decision Tree for WES samples (singletons)

For the interpretability reasons we put 2 decision trees here (same analysis as in the last chapter of the paper of in-house samples sequenced with ssHAEv6 and ssHAEv7 enrichment kits and also analysed with different CytoScape arrays) in fig. 16.

Features that were used for splits:

1. AverageLoglikPerTileCorrect – log-likelihood score, divided by the number of enrichment regions from the bed file, affected by CNV;
2. AverageNumOfMarkers – how many enrichment regions are inside the CNV;
3. AverageLoglicPerTile – log-likelihood score, divided by 1000 (score per length of variant in kbs);
4. AverageLoglikScore – raw score of variant;
5. AverageQval – q-value of the variant.

Worth to mention that, even if the log-likelihood score itself was not used in these trees, it is actually still detected as the one having the maximum importance via random forest approach (fig. 17).

| | FDR, sites | FDR, CNVs | # sites |
|---|---|---|---|
| ClinCNV del (34) | 0.098 | 0.002 | 150 |
| ClinCNV dup | 0.24 | 0.018 | 188 |

**Table 1:** *FDR of ClinCNV (filtered set, large cohort used for normalization)*


[Venables et al., 2002] Venables, W. N. and Ripley, B. D. (2002) Modern Applied Statistics with S. Fourth Edition. Springer, New York. ISBN 0-387-95457-0

[Fromer et al., 2012] Fromer, M., Moran, J. L., Chambert, K., Banks, E., Bergen, S. E., Ruderfer, D. M., ... and Purcell, S. M. (2012). Discovery and statistical genotyping of copy-number variation from whole-exome sequencing depth. *American journal of human genetics*, 91(4), 597–607. doi:10.1016/j.ajhg.2012.08.005

[Campbell et al., 2017] Campbell, P., Getz, G., Stuart, J., Korbel, J., Stein, L. (2017) Pancancer analysis of whole genomes *bioRxiv*, https://doi.org/10.1101/162784

[Whitworth et al., 2018] Whitworth, J., Smith, P. S., Martin, J. E., West, H., Luchetti, A., Rodger, F., ... Maher, E. R. (2018). Comprehensive Cancer-Predisposition Gene Testing in an Adult Multiple Primary Tumor Series Shows a Broad Range of Deleterious Variants and Atypical Tumor Phenotypes *American journal of human genetics*, 103(1), 3-18. doi:10.1016/j.ajhg.2018.04.013

[Puente et al., 2015] Puente, X.S., Bea S., Valdes-Mas, R., Villamor, N., Gutierrez-Abril. J., Martin-Subero, J.I., Munar, M., ..., Campo, E. (2015) Non-coding recurrent mutations in chronic lymphocytic leukaemia. *Nature.* 2015 Oct 22;526(7574):519-24. doi: 10.1038/nature14666. Epub 2015 Jul 22.


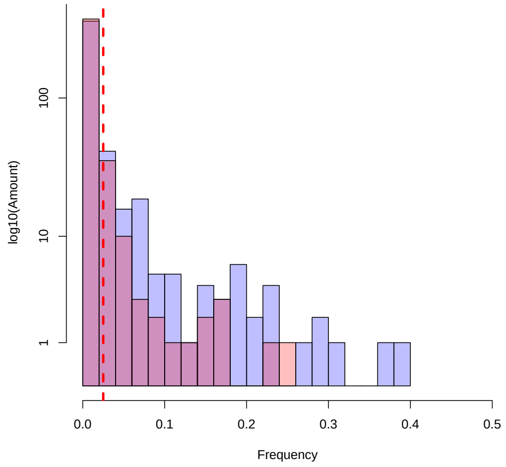


**(a)** *Frequency of CNV sites (v4, 5% FDR).*

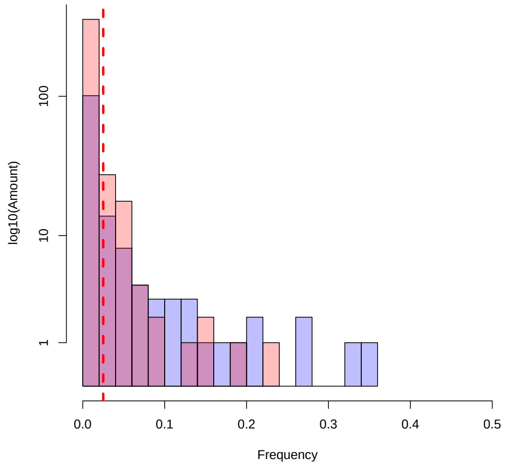


**(b)** *Frequency of CNV sites (v5, 12% FDR).*

**Figure 13:** *Overview of deletions and duplications frequencies in 2 WES callsets. Vertical line denotes 2.5% of allele frequency. Deletions are shown in red, duplications – in blue.*

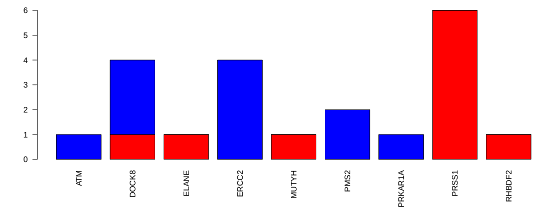


**(a)** *CNVs in cancer predisposition genes (v4 dataset, 5% FDR).*

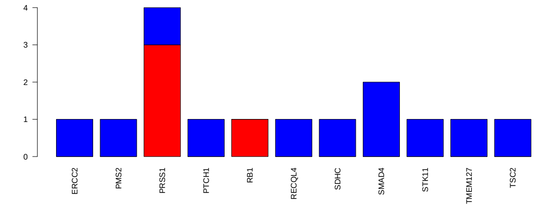


**(b)** *CNVs in cancer predisposition genes (v5 dataset, 12% FDR).*

**Figure 14:** *Overview of deletions and duplications in cancer predisposition genes in 2 WES callsets, affected by different CNVs: deletions (red) and duplications (blue).*

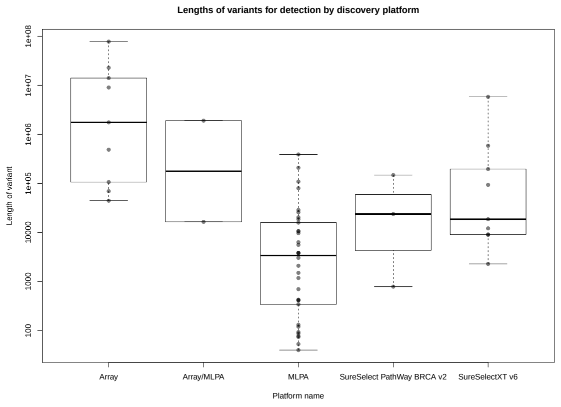


**Figure 15:** *Overview of CNVs for detection in shallow WGS data. Length of variant is shown on y-axis, platform used for detection of variant is shown on x-axis.*

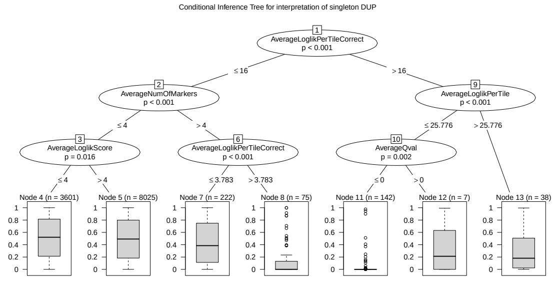


**(a)** *Decision tree for Duplications (singletons).*

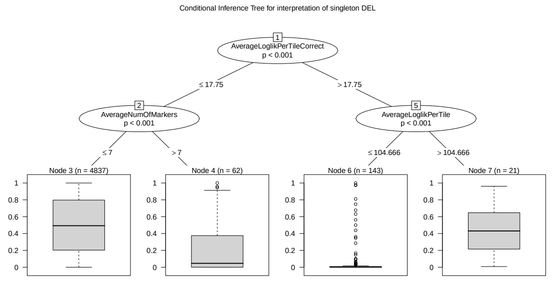


**(b)** *Decision tree for Deletions (singletons).*

**Figure 16:** *Decision trees for singleton CNVs in our cohort. Lower predicted value = higher chances of the variant to be real. FDR may be estimated as twice the number of values bigger than 0.5 in particular bin.*

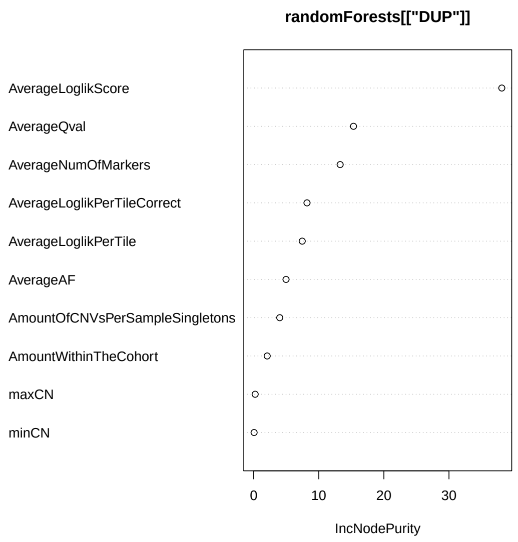


**(a)** *Variable importance for Duplications.*

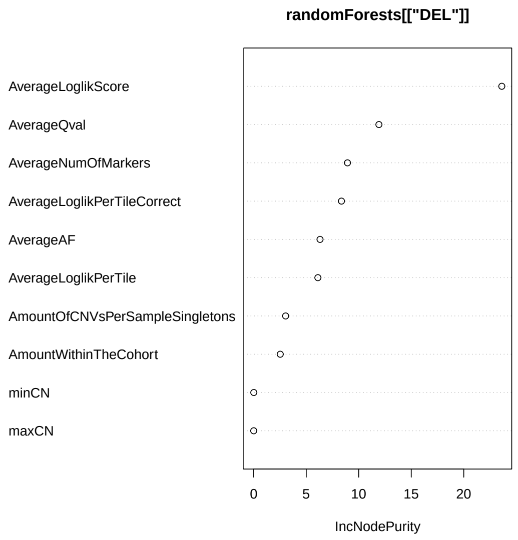


**(b)** *Variable importance for Deletions.*

**Figure 17:** *Variance importance of Random Forest FDR assignement approach.*
